# Conditional *in vivo* deletion of LYN kinase has little effect on a *BRCA1* loss-of-function-associated mammary tumour model

**DOI:** 10.1101/2023.03.23.533973

**Authors:** Giusy Tornillo, Lauren Warrington, Howard Kendrick, Adam T. Higgins, Trevor Hay, Sam Beck, Matthew J. Smalley

## Abstract

LYN kinase is expressed in BRCA1 loss-of-function-dependent mouse mammary tumours, in the cells of origin of such tumours, and in human breast cancer. Suppressing LYN kinase activity in BRCA1-defective cell lines, as well as in *in vitro* cultures of *Brca1*-null mouse mammary tumours, is deleterious to their growth. Here, we have examined the interaction between LYN kinase and BRCA1 loss-of-function in an *in vivo* mouse mammary tumour model, using conditional knockout *Brca1* and *Lyn* alleles. Comparison of *Brca1* tumour cohorts showed little difference in mammary tumour formation between animals that were wild type, heterozygous or homozygous for the conditional *Lyn* allele, although this was confounded by factors including incomplete *Lyn* recombination in some tumours. RNAseq analysis demonstrated that tumours with high levels of *Lyn* gene expression had a slower doubling time, but this was not correlated with levels of LYN staining in tumour cells themselves. Rather, high *Lyn* expression and slower tumour growth were likely a result of B-cell infiltration. The multifaceted role of LYN means it is likely to present difficulties as a therapeutic target in breast cancer.

## Introduction

The protein product of the *BRCA1* gene is well-established as a tumour suppressor, principally of breast and ovarian cancer (Fu et al., 2022; Molyneux et al., 2010). Women who inherit one functional and one mutated copy are at an approximately 70% lifetime risk of developing breast cancer and 40% lifetime risk of ovarian cancer (Kuchenbaecker et al., 2017). Tumour formation is associated with loss-of-heterozygosity events which delete the functional copy (Mahdavi et al., 2019) and is accelerated by concomitant inactivation of the TP53 tumour suppressor protein (Kim et al., 2016; Liu et al., 2007; Zhu et al., 2022). However, carriers of *BRCA1* mutations may also show functional haploinsufficiency which increases the risk of overt neoplasia (Lim et al., 2009).

The most well-characterised role of BRCA1 is as a key component of error-free, homologous recombination-dependent repair of double-stranded DNA damage (Foo and Xia, 2022; O’Donovan and Livingston, 2010). It is the defect in this process in BRCA1 loss-of-function associated cancer that is exploited by the use of PARP inhibitors for therapy (Mateo et al., 2019). However, BRCA1 has a number of additional functions not directly affecting DNA damage repair (although they may influence the process indirectly). These include acting as an E3 ubiquitin ligase, regulating transcription and control of centrosomal replication (Densham and Morris, 2017; Nolan et al., 2017; Yoshida and Miki, 2004).

The cells of origin of BRCA1-associated mammary tumours, the mammary luminal epithelial estrogen receptor negative stem/progenitor population, express the c-KIT receptor tyrosine kinase at high levels, as well as its downstream pathway member, the SRC-family kinase LYN (Regan et al., 2012; Tornillo et al., 2018). LYN is overexpressed in human Triple Negative Breast Cancer (TNBC; the breast cancer subtype most strongly associated with BRCA1 loss) (Choi et al., 2010; Croucher et al., 2013; Hochgrafe et al., 2010; Molyneux et al., 2010) and is also expressed at high levels in mammary tumours from *Brca1* conditional knockout mice (Molyneux et al., 2010). We have directly demonstrated using human breast cancer cell lines, primary cultures from *Brca1*-deleted mouse mammary tumours and cultures from human *BRCA1*-null patient-derived breast cancer xenografts, that BRCA1 loss results in activation of LYN and downstream pathways, including AKT, and a growth and survival advantage to mammary tumour cells (Tornillo et al., 2018). We therefore suggested that LYN is an oncogene in the context of BRCA1 loss, and a potential therapeutic target in BRCA1 loss-of-function breast and ovarian cancers. However, this has not yet been tested in a gold-standard *in vivo* knockout mouse model. Therefore, we obtained a conditional knockout LYN allele and crossed it to our established *BlgCre Brca1^f/f^ p53^+/-^* mouse mammary tumour model. Surprisingly, we found that in this system, *BlgCre Brca1^f/f^ p53^+/-^* mice carrying two conditional knockout *Lyn* alleles had a shorter overall survival than heterozygous or *Lyn* wild type *BlgCre Brca1^f/f^ p53^+/-^* mice, but no difference in mammary tumour specific survival. Tumours with low levels of *Lyn* gene expression grew faster than tumours with high levels of *Lyn*, however, the extent of LYN protein expression in tumour cells as assessed by immunohistochemistry was not correlated with tumour growth (although we were not able to assess LYN kinase activity in tumour cells). Rather, increasing abundance of B-cells in tumours was significantly associated with a slower tumour doubling time. As B-cells also express *Lyn*, this likely explained the correlation of tumour doubling time with *Lyn* expression levels but not LYN staining in tumour cells. Our results suggest that as a therapeutic target in breast cancer, LYN kinase is likely to present difficulties.

## Results

### The *Lyn^fl(ex4)^* allele is efficiently recombined *ex vivo* resulting in loss of *Lyn* expression

We previously assessed the relationship between LYN and (*BRCA1*-associated) mammary tumourigenesis using, among other approaches, shRNA knockdown with two independent shRNA sequences. We controlled for off-target effects by re-expressing a *Lyn* cDNA which was resistant to the effects of the knockdown. Our findings suggested that functional LYN kinase was required for survival of BRCA1-null mammary tumour cells (Tornillo et al., 2018). However, the role of LYN in *BRCA1*-associated mammary tumourigenesis has not been investigated using the gold standard of conditional *in vivo* knockouts. Therefore, we obtained a mouse line carrying a conditional (*floxed* exon 4) *Lyn* allele (the *Lyn^tm1c^*allele; hereafter *Lyn^fl^*) from the Mary Lyon Centre, Harwell **(Fig. 1A)**. Full details of these mice, the breeding strategies in which they were used to generate experimental cohorts and genotyping primers are provided in the **Materials and Methods**, **Fig. S1** and **Table S1**.

**Figure 1:**
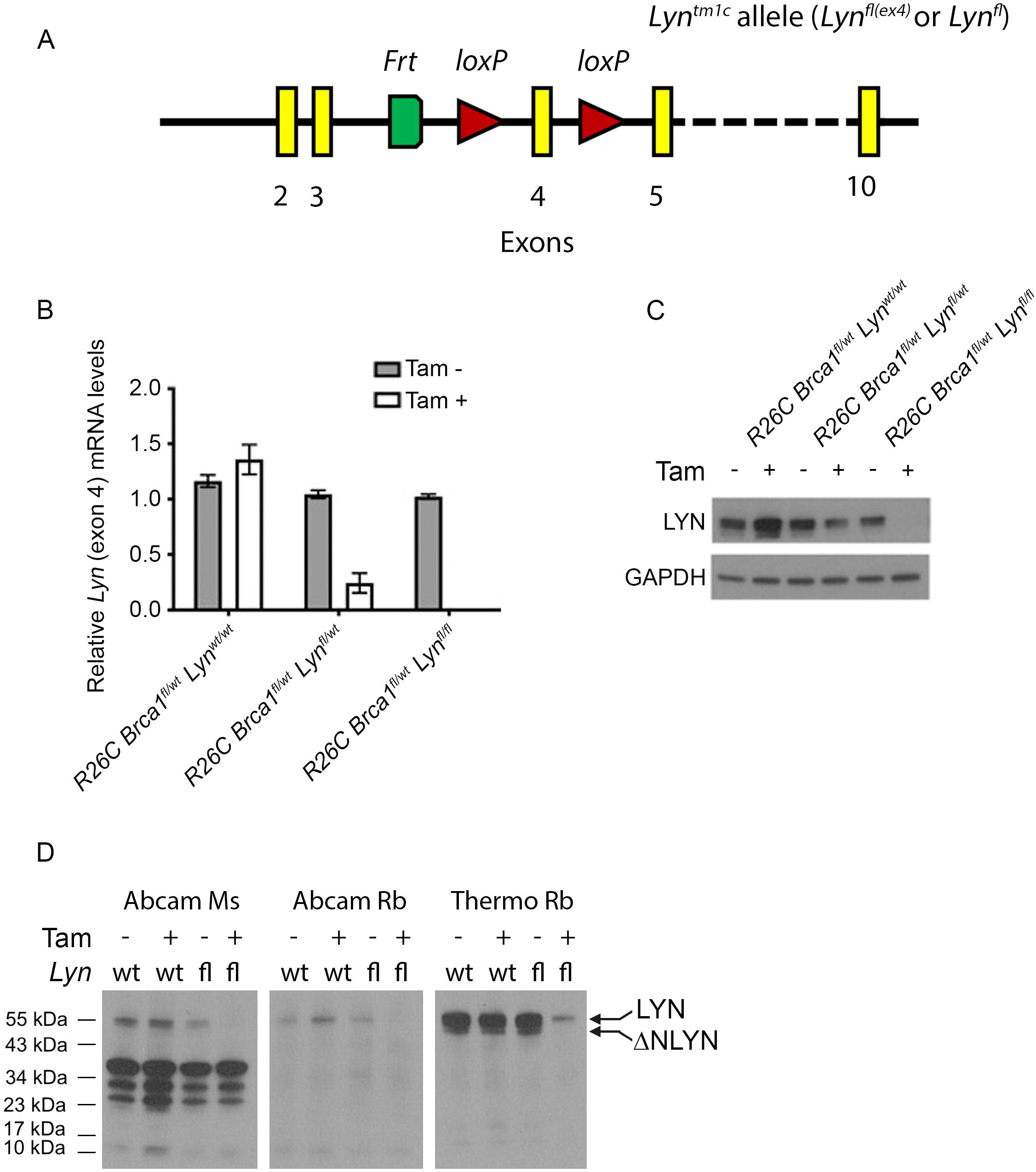
The *Lyn^fl^* allele is efficiently recombined *ex vivo* to deplete LYN protein. **A)** Schematic of the *Lyn^tm1c^* (*Lyn^fl(ex4)^* or *Lyn^fl^*) allele as supplied by the Mary Lyon Centre, MRC Harwell. Location of *loxP* sites flanking exon 4 indicated by red triangles. The single *Frt* site (green) is a remnant of the targeting strategy used to generate the allele. **B)** Quantitative real-time rtPCR analysis of *Lyn* levels using an exon 4-specific probe in cultured primary mouse mammary epithelial cells from *R26C Brca1^fl/+^ Lyn^wt/wt^*, *R26C Brca1^fl/+^ Lyn^fl/wt^*and *R26C Brca1^fl/+^ Lyn^fl/fl^* mice, treated with either vehicle control or tamoxifen. Results expressed as mean relative exon 4 levels ± 95% confidence intervals compared to to vehicle-treated *R26C Brca1^fl/+^ Lyn^fl/wt^* cells. **C)** Western blot for LYN expression in protein extracts from the cell cultures analysed in (B) (Thermofisher polyclonal antibody). **D)** Western blot analysis of LYN expression in *R26C Brca1^fl/+^ Lyn^wt/wt^* and *R26C Brca1^fl/+^ Lyn^fl/fl^* primary mammary epithelial cells, with or without tamoxifen treatment, using three different commercial anti-LYN antibodies. Each antibody was tested on identical loading of the same protein extracts. The expected LYN band and the putative ΔNLYN product visible with the Thermofisher rabbit polyclonal antibody are indicated.

We first used *Lyn^fl^* mice to establish a cohort of animals in which expression of a tamoxifen-inducible CRE recombinase was driven from the ubiquitously active *Rosa26* promoter (hereafter *R26C*). This cohort also included a conditional *floxed Brca1* allele (*Brca1^fl(ex22-24)^*; hereafter *Brca1^fl^*) (McCarthy et al., 2007; Molyneux et al., 2010). To test the recombination of the *Lyn^fl^* allele, mammary epithelial cells were harvested from *R26C Brca1^fl/wt^ Lyn^wt/wt^*, *R26C Brca1^fl/wt^ Lyn^fl/wt^*or *R26C Brca1^fl/wt^ Lyn^fl/fl^* mice and cultured as three-dimensional organoids according to our previous protocols (Tornillo et al., 2018). After one day, cultures were treated with 100 nM 4-hydroxytamoxifen (4OHT) or vehicle. After overnight incubation, cultures were washed to remove 4OHT and then cultured for a further 72 hours prior to lysis for isolation of either RNA or protein. Quantitative real-time rtPCR (qrtPCR) analysis (**Fig. 1B**) demonstrated no difference in expression of *Lyn exon 4* between vehicle-treated *R26C Brca1^fl/wt^ Lyn^wt/wt^*, *R26C Brca1^fl/wt^ Lyn^fl/wt^* or *R26C Brca1^fl/wt^ Lyn^fl/fl^* cells, or between vehicle and 4OHT-treated cells from *R26C Brca1^fl/wt^ Lyn^wt/wt^*mice. However, there was a significant reduction in *Lyn exon 4* levels in 4OHT-treated compared to vehicle-treated *R26C Brca1^fl/wt^ Lyn^fl/wt^* cells, while *Lyn exon 4* was undetectable in 4OHT-treated cells from *R26C Brca1^fl/wt^ Lyn^fl/fl^* mice. Western blot analysis of protein extracts from these cultures (**Fig. 1C**) confirmed these results.

To provide further evidence that deletion of *Lyn exon 4* results in a complete loss of LYN protein rather than, for example, generating a truncated protein which may have dominant negative effects, protein lysates from *R26C Brca1^fl/wt^ Lyn^wt/wt^* and *R26C Brca1^fl/wt^ Lyn^fl/fl^* cells, treated with either vehicle or 4OHT, were analysed by western blot using three different anti-LYN antibodies, one mouse monoclonal (Abcam ab1890) and two rabbit polyclonal (Abcam ab32398 and Thermofisher PA5-81925) in parallel (**Fig. 1D; Table S1**). In all other respects, the analysis was run identically, with identical amounts of protein loaded in all lanes. All three antibodies showed a substantial reduction in the amount of LYN protein detected in 4OHT-treated *R26C Brca1^fl/wt^ Lyn^fl/fl^* cells, compared to the other samples, and indeed in the mouse monoclonal and Abcam polyclonal samples LYN was undetectable. The mouse monoclonal antibody generated a number of non-specific bands only observed with the other reagents in very over-exposed blots (see **Supplemental Data File**). The Abcam polyclonal gave only a faint single band at the expected size which disappeared in 4OHT-treatment of *Lyn^fl^* cells. The Thermofisher rabbit polyclonal gave a strong signal of the expected size which was substantially reduced in the 4OHT-treated *R26C Brca1^fl/wt^ Lyn^fl/fl^* sample, although a faint band was still detectable, suggesting 100% recombination was not achieved in the cultures. The lower band visible with the Thermofisher polyclonal was not an artefact associated with the *Lyn^fl^* allele, as it was visible in extracts from both *Lyn^wt^*and *Lyn^fl^* cells. It was specific to *Lyn*, as it disappeared upon 4OHT-treatment of *Lyn^fl^*cells. It was not, however, the LYNB isoform (Tornillo et al., 2018), as the two LYN isoforms, LYNA and LYNB, are not fully resolved in the gradient gels used here (Tornillo et al., 2018). We suggest that the lower band is an endogenous product of caspase cleavage of LYN; a ΔNLYN variant has been previously described (Marchetti et al., 2009). Therefore, the *Lyn^fl^* allele is recombined by CRE recombinase and as a result is unable to generate protein.

### A *BlgCre Brca1^fl/fl^ p53^+/-^ Lyn^fl/fl^* mouse cohort has reduced overall survival but few other differences compared to cohorts with wild type *Lyn* alleles

Next, three cohorts of mice in which CRE expression was driven by the *Beta-lactoglobulin* promoter (*BlgCre*) were established. All three were homozygous for *floxed Brca1* alleles and also germline heterozygous for *p53* (*BlgCre Brca1^fl/fl^ p53^+/-^*), replicating the lines we have previously used (McCarthy et al., 2007; Molyneux et al., 2010). One cohort was wild type for *Lyn* (*BlgCre Brca1^fl/fl^ p53^+/-^ Lyn^wt/wt^*n=19), in the second animals were heterozygous for the conditional *Lyn* allele (*BlgCre Brca1^fl/fl^ p53^+/-^ Lyn^fl/wt^* n=22), in the third they were homozygous for the conditional *Lyn* allele (*BlgCre Brca1^fl/fl^ p53^+/-^ Lyn^fl/fl^* n=21). Mice were aged until defined humane endpoints were reached, at which point animals were euthanised and underwent a full necropsy. Where mice developed mammary tumours, these were regularly measured to determine tumour doubling times prior to the point at which euthanasia was necessary. Full details of all cohort animals and their pathology is provided in **Tables S2 and S3**.

We hypothesised that as LYN kinase activity was required for survival of cells that had lost BRCA1 activity (Tornillo et al., 2018), introducing the conditional *Lyn^fl(ex4)^* allele into the *BlgCre Brca1^fl/fl^ p53^+/-^* background would result in a significant increase in overall survival (i.e. age at which mice had to be euthanised for any reason) and also in mammary tumour-specific survival (i.e. the age at which mice had to be euthanised specifically as a result of the size of a mammary tumour). In contrast to our hypothesis, however, overall survival for *BlgCre Brca1^fl/fl^ p53^+/-^ Lyn^fl/fl^* mice was slightly, but significantly, shorter than for *BlgCre Brca1^fl/fl^ p53^+/-^ Lyn^fl/wt^* mice (median survival 350 days vs 365 days, respectively), although not significantly different to *BlgCre Brca1^fl/fl^ p53^+/-^ Lyn^wt/wt^* mice (median survival 366 days) (**Fig. 2A**). There was no significant difference in mammary tumour-specific survival between the cohorts (**Fig. 2B**).

**Figure 2:**
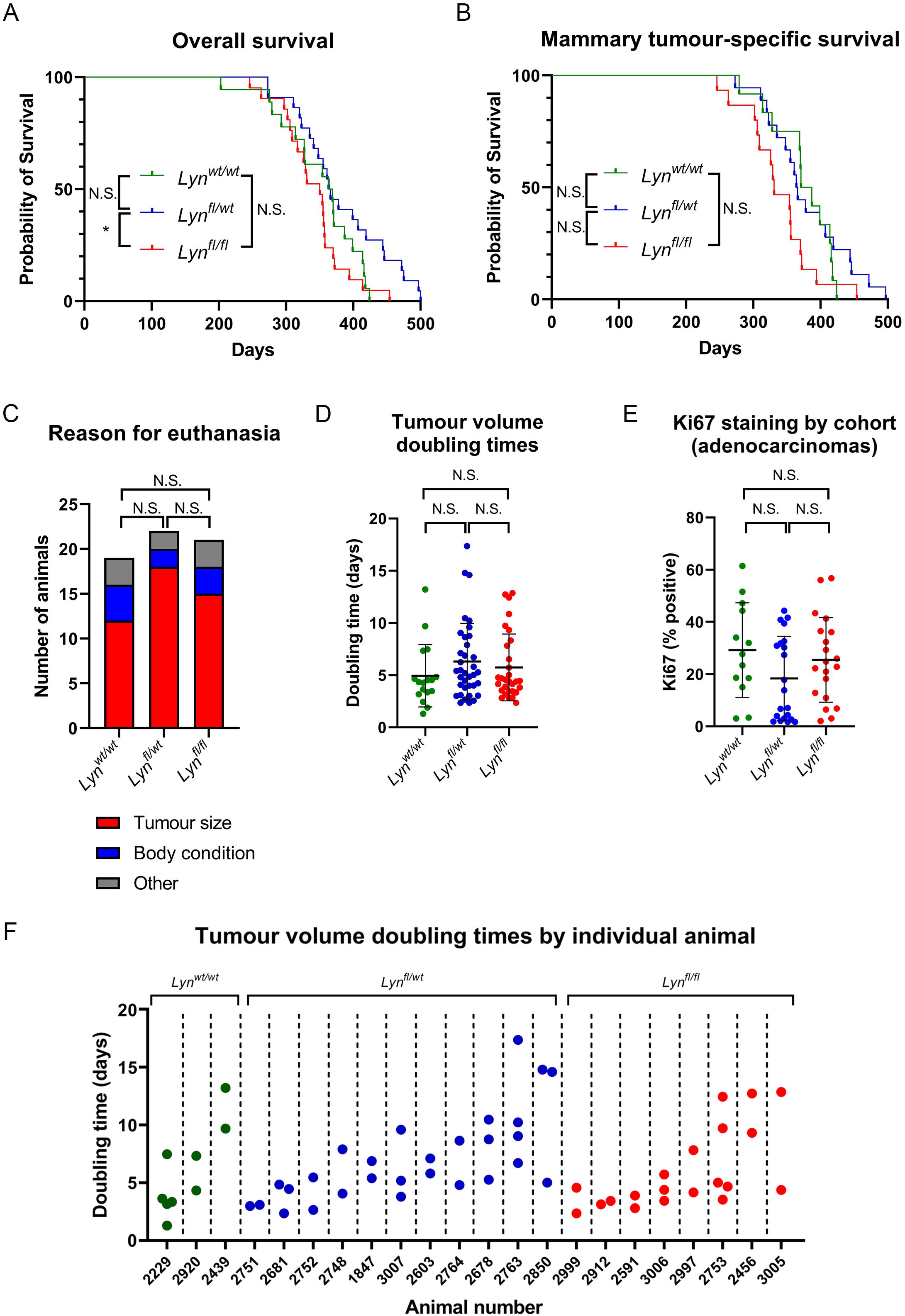
*Lyn^fl/fl^*mice have decreased overall survival in the *BlgCre Brca1^fl/fl^ p53^+/-^* model and have tumours with highly heterogeneous growth characteristics. **A)** Kaplan-Meier curve of overall survival of *BlgCre Brca1^fl/fl^ p53^+/-^ Lyn^wt/wt^* (n=19), *BlgCre Brca1^fl/fl^ p53^+/-^ Lyn^fl/wt^* (n=22) and *BlgCre Brca1^fl/fl^ p53^+/-^ Lyn^fl/fl^* (n=21) cohorts. *Lyn^fl/fl^*mice have a significantly shorter survival than *Lyn^fl/wt^* mice (P<0.05; Log Rank test). **B)** Kaplan-Meier curve of survival of *BlgCre Brca1^fl/fl^ p53^+/-^ Lyn^wt/wt^* (n=12), *BlgCre Brca1^fl/fl^ p53^+/-^ Lyn^fl/wt^* (n=18) and *BlgCre Brca1^fl/fl^ p53^+/-^ Lyn^fl/fl^* (n=15) cohorts considering only mice euthanased as a result of a mammary tumour reaching a specified endpoint. There are no significant differences (Log Rank test). **C)** Reason for euthanasia in *BlgCre Brca1^fl/fl^ p53^+/-^ Lyn^wt/wt^* (n=19), *BlgCre Brca1^fl/fl^ p53^+/-^ Lyn^fl/wt^* (n=22) and *BlgCre Brca1^fl/fl^ p53^+/-^ Lyn^fl/fl^* (n=21) cohorts. No significant differences (Chi^2^ test for trend). **D)** Doubling times (days) for individual tumours in each cohort (n=17, 36, 30 for *Lyn^wt/wt^*, *Lyn^fl/wt^* and *Lyn^fl/fl^* respectively). The value for each tumour is plotted with the mean±s.d.. No significant differences (one-way ANOVA across cohorts and also two-tailed t-tests comparing each cohort to the others). **E)** Ki67 percentage positivity in sections of mammary adenocarcinomas from each cohort (n=13, 21, 20 for *Lyn^wt/wt^*, *Lyn^fl/wt^*and *Lyn^fl/fl^* respectively). The value for each tumour is plotted with the mean±s.d.. No significant differences (Kruskel-Wallis across cohorts and Mann-Whitney tests comparing each cohort to the others). **F)** Tumour volume doubling times (days) by animal showing only animals from each cohort with more than one tumour measured.

The majority of mice in all cohorts were euthanised because of the growth of the mammary tumours (**Fig. 2C**). Other reasons for euthanasia included scratches, vestibular syndrome or poor body condition score, with a number of examples of non-mammary neoplasia found upon necropsy, as well as reactive hyperplasia of the spleen. In seven cases, neoplastic epithelial deposits were observed in the lungs of animals carrying mammary tumours (**Fig. S2A,B**); in one of these cases, both the lung deposit and the primary tumour had a squamous histology, consistent with metastatic spread of the primary tumour (**Fig. S2B**). Other neoplastic lesions included haemangiosarcoma (one case) and osteosarcoma (two cases) (**Fig. S2C-F**). Histological analysis of enlarged spleens suggested this was largely reactive but in some cases the histology was consistent with histiocytic sarcoma (two cases; not shown) or lymphoma (clonal analysis of B- and/or T-cell receptor rearrangement to confirm lymphoma was not carried out as this was not the primary focus of this study). Florid extra-medullary haematopoiesis was also noted in some cases. There was no visible difference in LYN staining in the white pulp of the spleen in *Lyn^fl/fl^* animals compared to others, with an expected decrease in LYN staining in proliferating B-cell germinal centres (**Fig. S3**).

There were no significant differences in histotype of mammary tumours between the cohorts (the majority of which were adenocarcinomas of no special type; **Figs. S4 and S5A**). Many animals developed more than one tumour, and six of these developed more than one tumour in the same gland (**Supplementary Table 2**), but there was no significant difference between the cohorts in terms of numbers of tumours developed by each animal (**Fig. S5B**). The growth of every mammary tumour that was palpable while an animal was alive was measured daily until a humane endpoint was reached. This enabled doubling times for every tumour with three or more measurements to be established. There was no significant difference in doubling times of mammary tumours between the cohorts (**Fig. 2D**). This was confirmed by Ki67 staining of sections from mammary tumours across the cohorts (considering only adenocarcinomas to eliminate different tumour histotypes as a potential confounding factor) (**Fig. 2E; Table S4**). Within each cohort there was, however, considerable heterogeneity in tumour doubling times and when animals which developed more than one tumour were considered individually, it was apparent that even in a single animal, doubling times of tumours could vary widely (**Fig. 2F**).

### Cohort genotype only partly predicts LYN expression in tumours

Variation in behaviour of tumours across a cohort, and indeed in multiple tumours from a single mouse, could result from partial *floxed* allele recombination in *Lyn^fl/wt^* or *Lyn^fl/fl^* mice, or from suppression of LYN expression by other mechanisms in *Lyn^wt/wt^* mice. If such variation existed, it would confound any analysis of the role of LYN in *Brca1*-dependent mammary tumourigenesis based solely on cohort genotype.

Therefore, to directly assess LYN expression levels in tumours from the three cohorts, thirteen *Lyn^wt/wt^* tumours, twenty-one *Lyn^fl/wt^* tumours and twenty *Lyn^fl/fl^* tumours, all adenocarcinomas (no special type) were randomly selected for staining for LYN protein (**Fig. 3**). LYN staining was assessed both qualitatively and semi-quantitatively using a histoscore approach based on strength of staining and the area of the tumour stained (see **Materials and Methods**, **Fig. S5C, Table S4**). Scoring was carried out blinded to genotype; once scored, tumours were unblinded and analysed.

**Figure 3:**
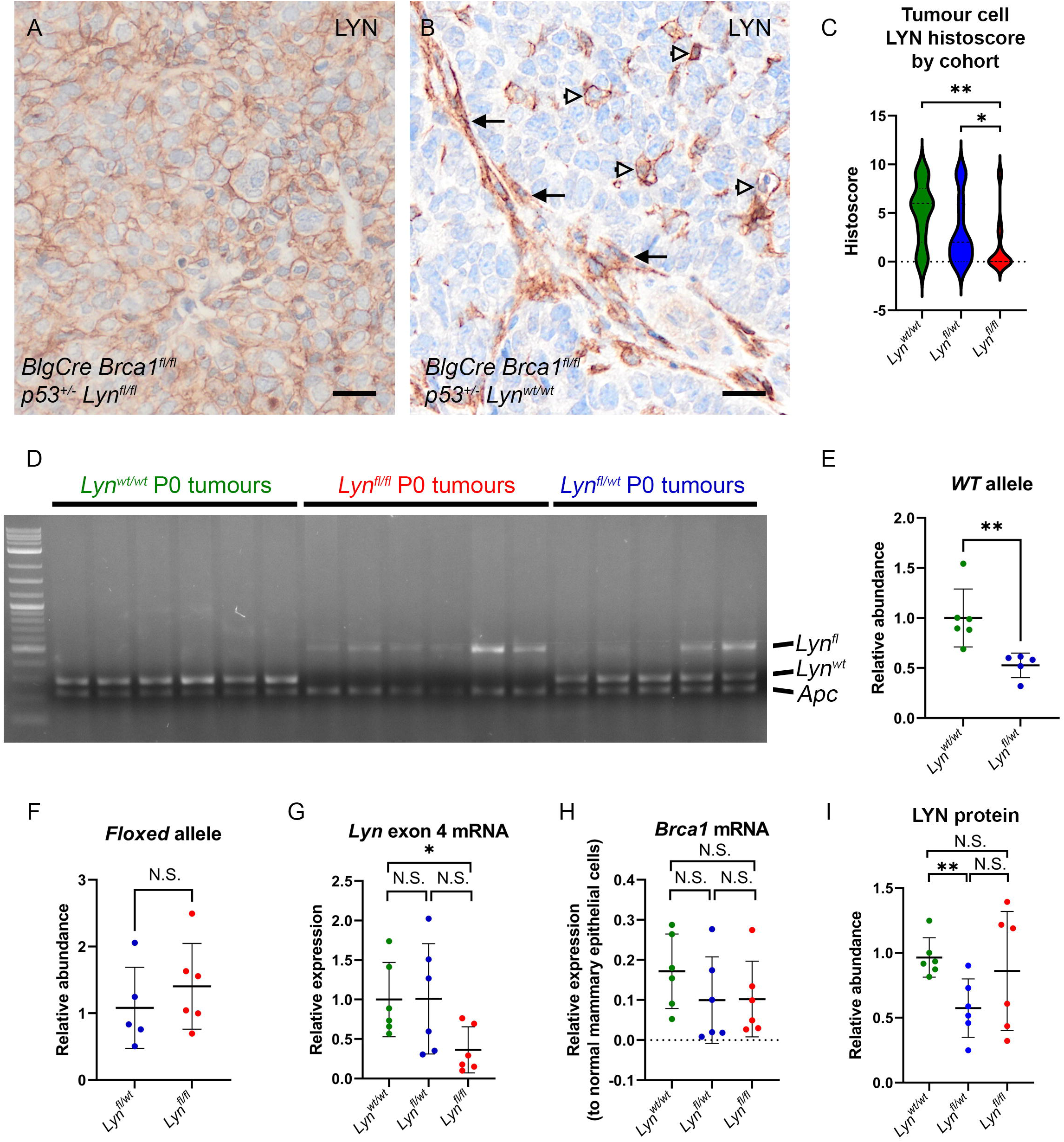
LYN expression in tumours is heterogeneous but decreased overall in *BlgCre Brca1^fl/fl^ p53^+/-^ Lyn^fl/fl^* mice. **A)** LYN staining pattern in epithelial-like tumour cells. Bar = 20 μm. **B)** LYN staining in single cells (white arrowheads) and cancer-associated fibroblast-like cells (black arrows). Bar = 20 μm. **C)** ‘Histoscore’ quantitation of LYN staining in epithelial-like tumour cells (see Methods for details; n=13, 21, 20 for *Lyn^wt/wt^*, *Lyn^fl/wt^* and *Lyn^fl/fl^* respectively). The *Lyn^fl/fl^*cohort has significantly less staining than the *Lyn^wt/wt^* or *Lyn^fl/wt^* cohort (**P<0.01, *P<0.05, Mann-Whitney test). **D)** Semi-quantitative PCR analysis of *Lyn^fl(ex4)^* and *Lyn^wt^* alleles in genomic DNA isolated from primary cultures of *BlgCre Brca1^fl/fl^ p53^+/-^ Lyn^wt/wt^*(n=6), *BlgCre Brca1^fl/fl^ p53^+/-^ Lyn^fl/wt^*(n=5) and *BlgCre Brca1^fl/fl^ p53^+/-^ Lyn^fl/fl^*(n=6) tumour cells. Analysis of the *Apc* locus was included as a control to enable relative quantitation. Each PCR reaction included the primers for all three alleles. **E)** Quantitation of relative abundance of *Lyn^wt^* alleles in (D), considering only *Lyn^wt/wt^* and *Lyn^fl/wt^*cultures (as no *Lyn^wt^* alleles are present in *Lyn^fl/fl^*cells). Data presented as abundance in each sample relative to the *Apc* band in that sample. The mean abundance±s.d. of each group is indicated. There are 50% fewer (P<0.01, Mann-Whitney test) *Lyn^wt^* alleles in the *Lyn^fl/wt^* compared to the *Lyn^wt/wt^* cultures, as expected. **F)** Quantitation of relative abundance of *Lyn^fl(ex4)^*alleles in (**D**), considering only *Lyn^fl/wt^* and *Lyn^fl/fl^*cultures (as no *Lyn^fl^* alleles are present in *Lyn^wt/wt^*cells). Data presented as abundance in each sample relative to the *Apc* band in that sample. The mean abundance±s.d. of each group is indicated. No significant difference between the samples, but the heterogeneity of the samples reflects the clear differences in band intensities seen in (**D**). **G)** Quantitative real-time rtPCR analysis of *Lyn exon 4* expression in primary tumour cell cultures. Data presented as expression relative to the mean value for the *Lyn^wt/wt^* cells (*P<0.05, Mann-Whitney test). **H)** Quantitative real-time rtPCR analysis of *Brca1* expression in primary tumour cell cultures. Data presented as expression relative to *Brca1* levels in lysates of freshly isolated normal mouse mammary epithelial cells for *Brca1* (no significant differences, Mann-Whitney tests). **I)** Relative expression of LYN protein in primary tumour cultures as determined by Western blot analysis, quantified relative to standard loading controls and normalised to one *Lyn^wt/wt^* tumour culture sample (see **Supplementary Data File** for raw blots). For **G-I**, n=6 samples of each genotype.

Staining patterns fell into three types: sheets and nests of epithelial-like tumour cells showing membrane staining (**Fig. 3A**); single cells scattered throughout the tumour which were typically cells with pseudopodia and an appearance suggest a motile phenotype (**Fig. 3B**, white arrowheads); cells with the appearance of tumour-associated fibroblasts (**Fig. 3B**, black arrows).

Histoscore quantitation of LYN staining was carried out for the neoplastic epithelial-like tumour cells. LYN staining of these varied significantly with tumour genotype (Kruskal-Wallis test, P=0.0063) (**Fig. 3C**). There was a significant reduction in LYN staining in the cells of *Lyn^fl/fl^* tumours compared to *Lyn^wt/wt^* tumours (Mann-Whitney test, P=0.002) and *Lyn^fl/wt^* tumours (Mann-Whitney, P=0.0344). However, some *Lyn^fl/fl^*tumours clearly retained strong LYN staining while some *Lyn^wt/wt^*tumours showed very little or no staining. LYN staining in *Lyn^fl/wt^*tumours was reduced compared to *Lyn^wt/wt^* but the difference was not statistically significant.

While these results showed that, overall, there was a correlation between LYN staining of a tumour and the genotype of the animal the tumour came from, they also highlighted the variability in staining between tumours of the same genotype. This suggested that genotype could not be fully relied upon to predict LYN expression in any one individual tumour. To understand more objectively the relationship between tumour genotype and LYN expression in tumour cells, in the absence of cells from the tumour microenvironment which may also express LYN, we isolated live cells from 6 tumours of each cohort (*Lyn^wt/wt^*, *Lyn^fl/wt^* and *Lyn^fl/fl^*) and cultured them in conditions optimised for epithelial tumour cell primary culture before harvesting DNA, RNA and protein for analysis (DNA was only available for analysis from five of the six *Lyn^fl/wt^* samples).

Semi-quantitative PCR analysis of the *Lyn^fl^* and *Lyn^wt^* alleles demonstrated that the abundance of the *Lyn^wt^*allele in primary cultures of *BlgCre Brca1^fl/fl^ p53^+/-^ Lyn^fl/wt^* tumours was approximately half that in cultures of *BlgCre Brca1^fl/fl^ p53^+/-^ Lyn^wt/wt^* tumours, as expected (**Fig. 3D,E**). However, there was also no significant difference overall in the abundance of the *Lyn^fl^* allele between *BlgCre Brca1^fl/fl^ p53^+/-^ Lyn^fl/wt^* and *BlgCre Brca1^fl/fl^ p53^+/-^ Lyn^fl/fl^* tumour cells (**Fig. 3D,F**), and it is clear that while in some tumours the *Lyn^fl^* allele had recombined effectively, in others it remained intact (**Fig. 3D**). Assessment of *Lyn* expression by qrtPCR using a probe targeting exon 4, demonstrated that there was a significant reduction in *Lyn* expression in *BlgCre Brca1^fl/fl^ p53^+/-^ Lyn^fl/fl^* tumour cells compared to wild-type cells, but again in some individual tumours *Lyn* exon 4 expression was comparable to that seen in *Lyn^wt/wt^*tumours (**Fig. 3G**). *Brca1* expression was, as expected very low in tumours from all three lines relative to normal mammary epithelial cells (**Fig. 3H**), although not zero, likely due to the presence of non-transformed, non-recombined epithelial cells derived from normal ducts trapped within the tumour and thus ‘contaminating’ the primary tumour cultures. Finally, assessment of LYN protein levels in these cells by western blot showed that *Lyn^fl/wt^*tumour cells had significantly less protein than *Lyn^wt/wt^* tumour cells, but again while some *Lyn^fl/fl^* tumour cells had low LYN protein levels, others had levels of LYN comparable to the wild type cultures (**Fig. 3I**).

### Transcriptional analysis of tumours demonstrates tumours with high *Lyn* expression have a slower doubling time

Analysis of LYN protein levels in tumours demonstrated that while overall there was a correlation between cohort genotype and LYN expression, there were a number of cases in which tumours from a *Lyn* wildtype mouse had very low or undetectable levels of LYN, while tumours from mice homozygous for the *Lyn* flox allele could actually have high levels of LYN expression. This, together with the presence of multiple tumours with different growth rates in some animals, confounded the analysis of the cohorts (**Fig. 2A,B**).

Therefore, to directly assess differences in the biology of the tumours from the cohorts, and to determine whether or not such differences were correlated with *Lyn* expression, we carried out an RNAseq analysis of tumour pieces from 39 tumours – 12 from *BlgCre Brca1^fl/fl^ p53^+/-^ Lyn^wt/wt^* mice, 14 from *BlgCre Brca1^fl/fl^ p53^+/-^ Lyn^fl/wt^* mice and 13 from *BlgCre Brca1^fl/fl^ p53^+/-^ Lyn^fl/fl^* mice. All were adenocarcinomas of no special type, to eliminate histological variation as a confounding factor. The sample details are provided in **Table S5**. The data were analysed in three ways, two of which were ‘supervised’ and one which was ‘unsupervised’. First, Differentially Expressed Genes (DEGs) significantly (<0.05 adjusted P-value; ≤0.05 or ≥2.0 log2 fold change) differentially expressed between tumours from *BlgCre Brca1^fl/fl^ p53^+/-^ Lyn^wt/wt^* and *BlgCre Brca1^fl/fl^ p53^+/-^ Lyn^fl/fl^* mice were identified (supervised analysis on the basis of genotype). Second, the normalised expression values for *Lyn* were used to rank the whole 39-sample tumour set from the strongest *Lyn* expressing to the weakest *Lyn* expressing tumour. Then, the 13 tumours most strongly expressing *Lyn* (‘*Lyn* high’ group) were compared to the 13 tumours with the weakest *Lyn* expression (‘*Lyn* low’ group) to identify DEGs (supervised analysis on the basis of *Lyn* expression). Finally, a Principal Component Analysis (PCA) analysis was carried out on the complete normalised data set of 39 tumours to identify any groups of tumours which could be distinguished from each other on the basis of transcriptional profiles in an unbiased manner. Significant DEGs were identified between the PCA groups (unsupervised analysis). The raw and normalised data for these comparisons are provided in **Tables S6 – S8**. Significant DEGs are provided in **Table S9**, Gene Set Enrichment Analysis (GSEA) using g:Profiler in **Table S10** and a summary of enriched Gene Ontogeny Bioprocess and KEGG pathways in **Table S11**.

With the *BlgCre Brca1^fl/fl^ p53^+/-^ Lyn^wt/wt^* and *BlgCre Brca1^fl/fl^ p53^+/-^ Lyn^fl/fl^* comparison, only 93 significant DEGs were identified, 12 upregulated in *Lyn^fl/fl^* tumours relative to *Lyn^wt/wt^* tumours and 81 downregulated in *Lyn^fl/fl^* tumours relative to *Lyn^wt/wt^* tumours (**Fig. 4A**; **Table S9**). This emphasised our previous findings that animal genotype was not necessarily a good surrogate for *Lyn* expression or differences in tumour biology. Indeed, there was no difference in *Lyn* expression, as defined by normalised RNAseq *Lyn* counts, between the cohorts (**Fig. 4B**).

**Figure 4:**
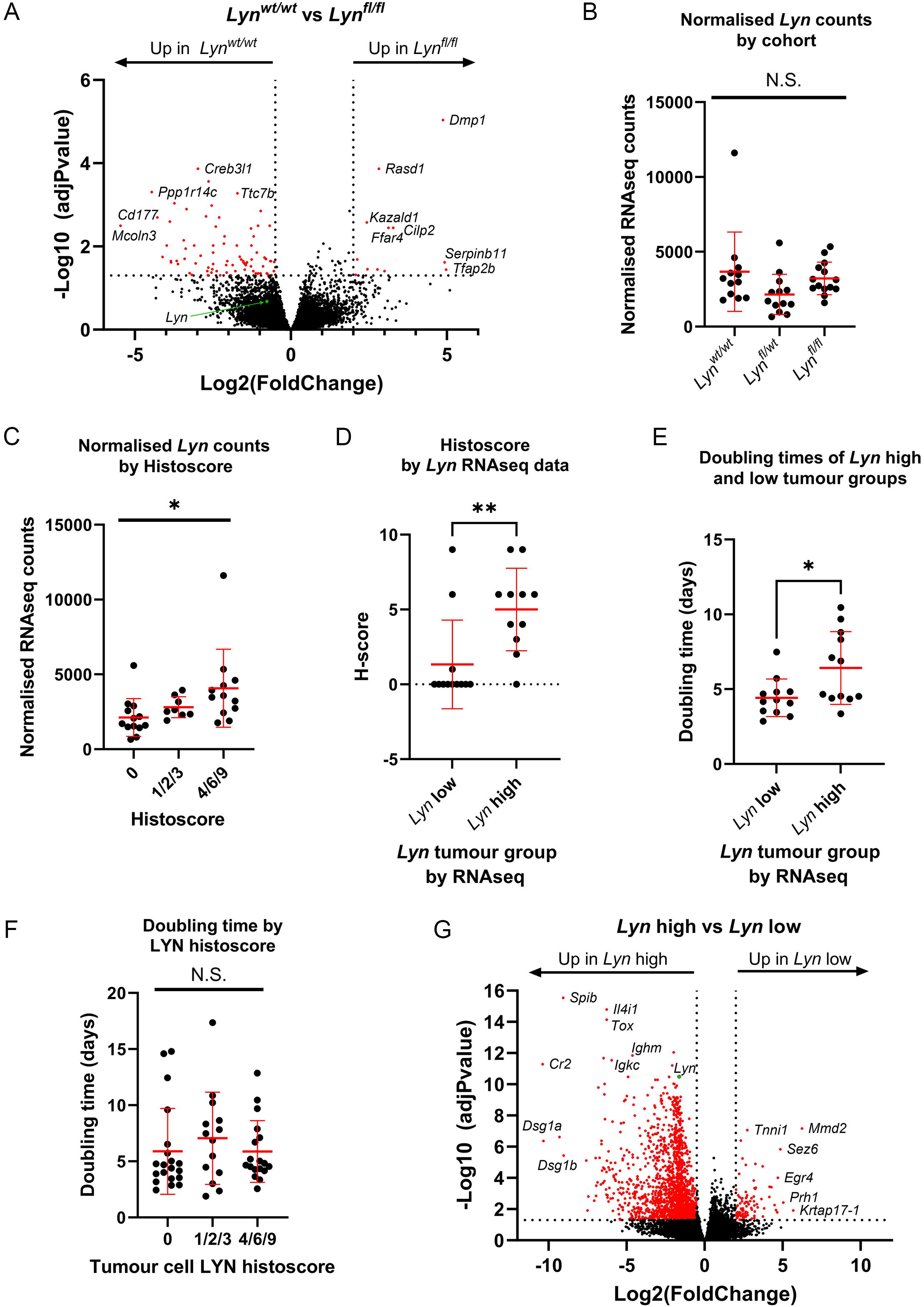
Tumour molecular profiles correlate with *Lyn* expression but not with tumour cohort. **A)** Volcano plot (-Log10(adjusted P value) against Log2(Fold change)) of differentially expressed genes comparing *Lyn^wt/wt^* (n=12) and *Lyn^fl/fl^*(n=14) tumours. Genes with an adjusted P value of <0.05 and a fold change of ≤0.5 or ≥2 were considered significant and are indicated in red. The most strongly differentially expressed genes are labelled. *Lyn* is indicated with a green dot and labelled. **B)** Normalised RNAseq *Lyn* counts in tumours from each cohort (n=12, 13,14 for *Lyn^wt/wt^*, *Lyn^fl/wt^* and *Lyn^fl/fl^* respectively). Mean±s.d.. No significant differences (ANOVA). **C)** Normalised RNAseq *Lyn* counts in tumours with different LYN histoscore grading (0, no LYN staining; 1/2/3, low LYN staining or strong staining but only in a small region; 4/6/9, moderate to strong LYN staining) (n=13, 8, 12, respectively). Increased *Lyn* counts are associated with increasing histoscore (ANOVA, *P<0.05). **D)** Histoscore of top tertile (*Lyn* high; n=12) vs bottom tertile (*Lyn* low; n=11) *Lyn* expressing tumours by RNAseq. Mean±s.d.. **P<0.01 (Mann-Whitney). **E)** *In vivo* doubling time (days) of top tertile (*Lyn* high; n=12) vs bottom tertile (*Lyn* low; n=12) *Lyn* expressing tumours by RNAseq. Mean±s.d.. *P<0.05 (t-test). **F)** *In vivo* doubling time (days) of tumours from LYN histoscore groups (n=20, 14, 18, for groups 0, 1/2/3, 4/6/9 respectively). Mean±s.d.. No significant differences (ANOVA). **G)** Volcano plot (-Log10(adjusted P value) against Log2(Fold change)) of differentially expressed genes comparing top tertile (*Lyn* high; n=13) vs bottom tertile (*Lyn* low; n=13) *Lyn* expressing tumours defined by RNAseq. Genes with an adjusted P value of <0.05 and a fold change of ≤0.5 or ≥2 were considered significant and are indicated in red. The most strongly differentially expressed genes are labelled. *Lyn* is indicated with a green dot and labelled.

Next, we examined *Lyn* expression in all tumours as determined by the RNAseq data (ignoring genotypes) and compared this to LYN staining (**Table S4**). Normalised *Lyn* counts were elevated in tumours with a LYN histoscore of 1/2/3 compared to tumours with a score of 0, and elevated further in 4/6/9 histoscore tumours compared to 1/2/3 scored tumours (P<0.05; ANOVA; **Fig. 4C**). Consistent with this, when the tumours defined by normalised *Lyn* counts in the RNAseq as ‘*Lyn* high’ (top 13 most strongly *Lyn* expression tumours) and ‘*Lyn* low’ tumours (13 tumours with the weakest *Lyn* expression) (**Table S5**) were compared, the histoscore of the *Lyn* high tumours was significantly greater than that of the *Lyn* low tumours (P<0.01; Mann-Whitney; **Fig. 4D**). There were, however, outliers showing that in some tumours there was not a direct correlation between *Lyn* expression by RNAseq and LYN IHC staining. We next compared the *in vivo* doubling time of the tumours defined as *Lyn* high and *Lyn* low (**Fig. 4E**) by RNAseq. *Lyn* high tumours had a significantly longer doubling time (P<0.05; t-test), suggesting that in general they grew more slowly than *Lyn* low tumours and that they formed a distinct biological group. However, when the same tumours were divided into groups based on LYN histoscore of tumour cells, as directly assessed by IHC, there were no significant differences between tumours with no, moderate or high levels of LYN staining (**Fig. 4F**). Therefore, tumour doubling time was correlated with *Lyn* expression in the tumours as a whole but not directly with LYN expression in the neoplastic cells.

There were 1655 significant DEGs upregulated in *Lyn* high relative to *Lyn* low tumours and 100 significant DEGs downregulated in *Lyn* high relative to *Lyn* low tumours (equivalent to 100 DEGs significantly upregulated in *Lyn* low relative to *Lyn* high tumours; **Fig. 4G**; **Table S9**). This number of DEGs, compared to the number identified when comparing by genotype, showed that categorising tumours by *Lyn* expression was better at defining sets of tumours with biologically meaningful differences than categorising tumours by the genotype of the cohort from which they were derived.

Finally, we used PCA analysis on the normalised RNAseq expression values for the whole tumour set to identify in an unbiased manner groups of tumours with similar gene expression patterns. This analysis initially suggested the tumours could be split into either four (PCA groups 1, 2, 3 and 4; **Fig. 5A**) or two groups (combined groups 1/2 and 3/4). The normalised *Lyn* counts, LYN Histoscore and *in vivo* tumour doubling time were compared across either the four PCA groups individually or combined into two groups (**Fig. 5** and **Fig. S5D - F**). Groups 3 and 4 had significantly elevated *Lyn* counts and a significantly higher LYN Histoscore than groups 1 and 2 (**Fig. S5D,E**); the differences were more marked when comparing groups 3 and 4 combined as a single group against groups 1/2 (**Fig. 5B,C**). The combined group 3/4 had a significantly slower *in vivo* doubling time than group 1/2 (**Fig. 5D**) but these differences were not significant in the four-group analysis (**Fig. 6F**). As the group 3 and 4 tumours appeared to behave similarly to each other, and the group 1 and 2 tumours also behaved similarly, the differences between the groups were more marked in the two-group analysis, the difference in doubling time suggested a real biological difference between groups 3/4 and 1/2, and the two-group approach would enable greater numbers of tumours to be compared in each group, we concentrated on the two-group approach and identified significant DEGs from PCA group 3/4 compared to PCA group 1/2. There were 835 significant DEGs upregulated in PCA group 3/4 relative to PCA group 1/2 and 2437 significant DEGs downregulated in PCA group 3/4 relative to PCA group 1/2 (equivalent to 2437 DEGs significantly upregulated in PCA group 1/2 relative to PCA group 3/4; **Fig. 5E**).

**Figure 5:**
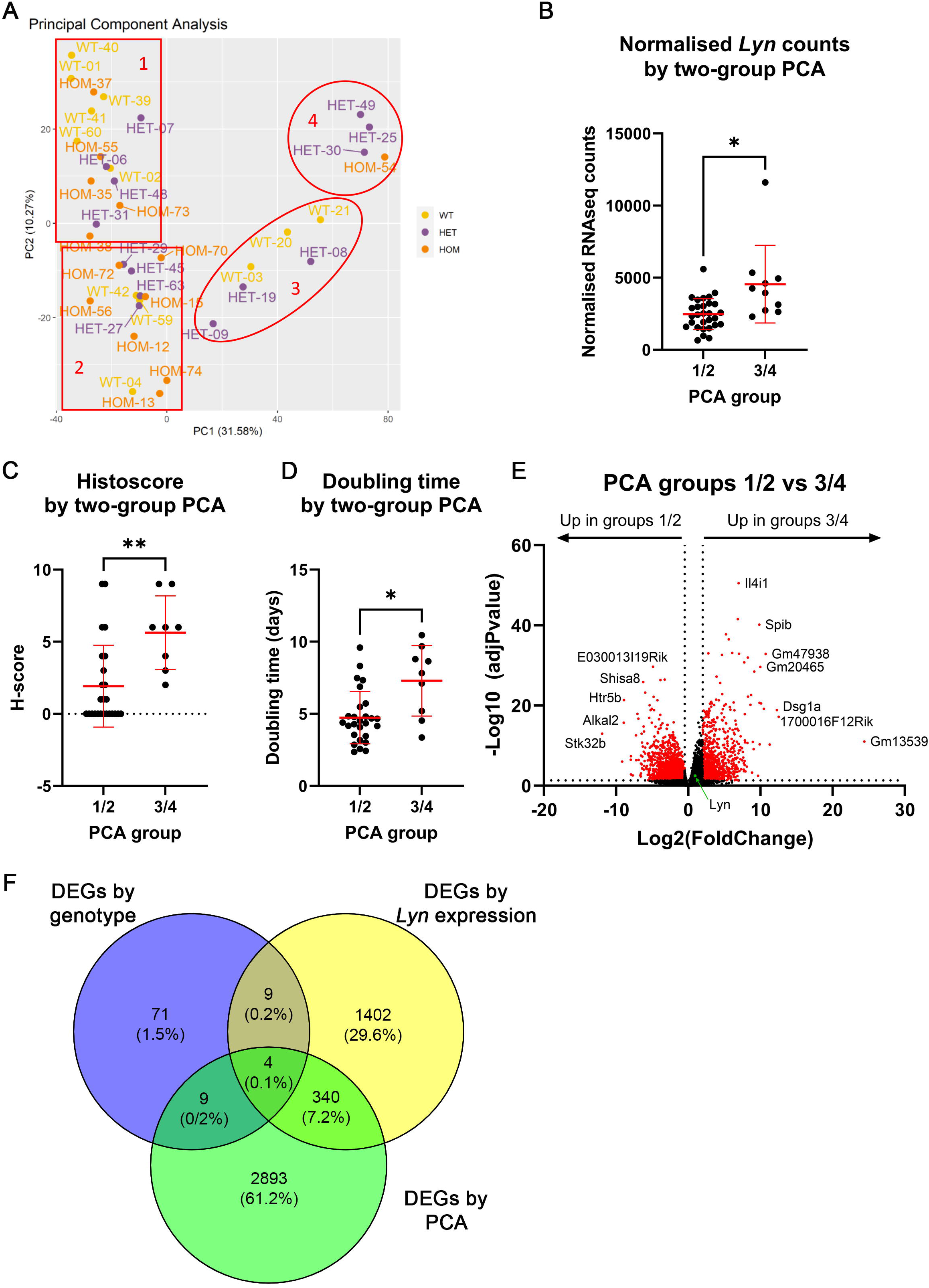
Principal Component Analysis (PCA) identifies two main groups of tumours which partially overlap with *Lyn* high and *Lyn* low tumours. **A)** PCA plot of 39 tumours analysed by RNAseq showing an unbiased assessment of tumour gene expression differences and similarities. PC1 divides the tumours into groups 1/2 (n=29) and 3/4 (n=10), PC2 further divides the tumours to give four groups; however, the majority of the differences between the tumour groups was generated by PC1. **B)** Normalised RNAseq *Lyn* counts in tumours from PCA groups 1/2 (n=29) and 3/4 (n=10). Mean±s.d.. *P<0.05 (two-tailed t-test). **C)** Histoscore of tumours from PCA groups 1/2 and 3/4. Mean±s.d.. **P<0.01 (Mann-Whitney). D) *In vivo* doubling time (days) of tumours from PCA groups 1/2 (n=27) and 3/4 (n=9). Mean±s.d.. *P<0.05 (t-test). **E)** Volcano plot (-Log10(adjusted P value) against Log2(Fold change)) of differentially expressed genes comparing tumours from PCA groups 1/2 (n=29) and 3/4 (n=10). Genes with an adjusted P value of <0.05 and a fold change of ≤0.5 or ≥2 were considered significant and are indicated in red. The most strongly differentially expressed genes are labelled. *Lyn* is indicated with a green dot and labelled. **F)** Venn diagram showing overlap between differentially expressed genes identified when comparing *Lyn^wt/wt^* vs *Lyn^fl/fl^* tumours, *Lyn* high vs *Lyn* low tumours and PCA group 1/2 vs 3/4 tumours. Note 340 genes overlap between the latter two groups but there is very little overlap with the *Lyn^wt/wt^* vs *Lyn^fl/fl^*tumour data.

**Figure 6:**
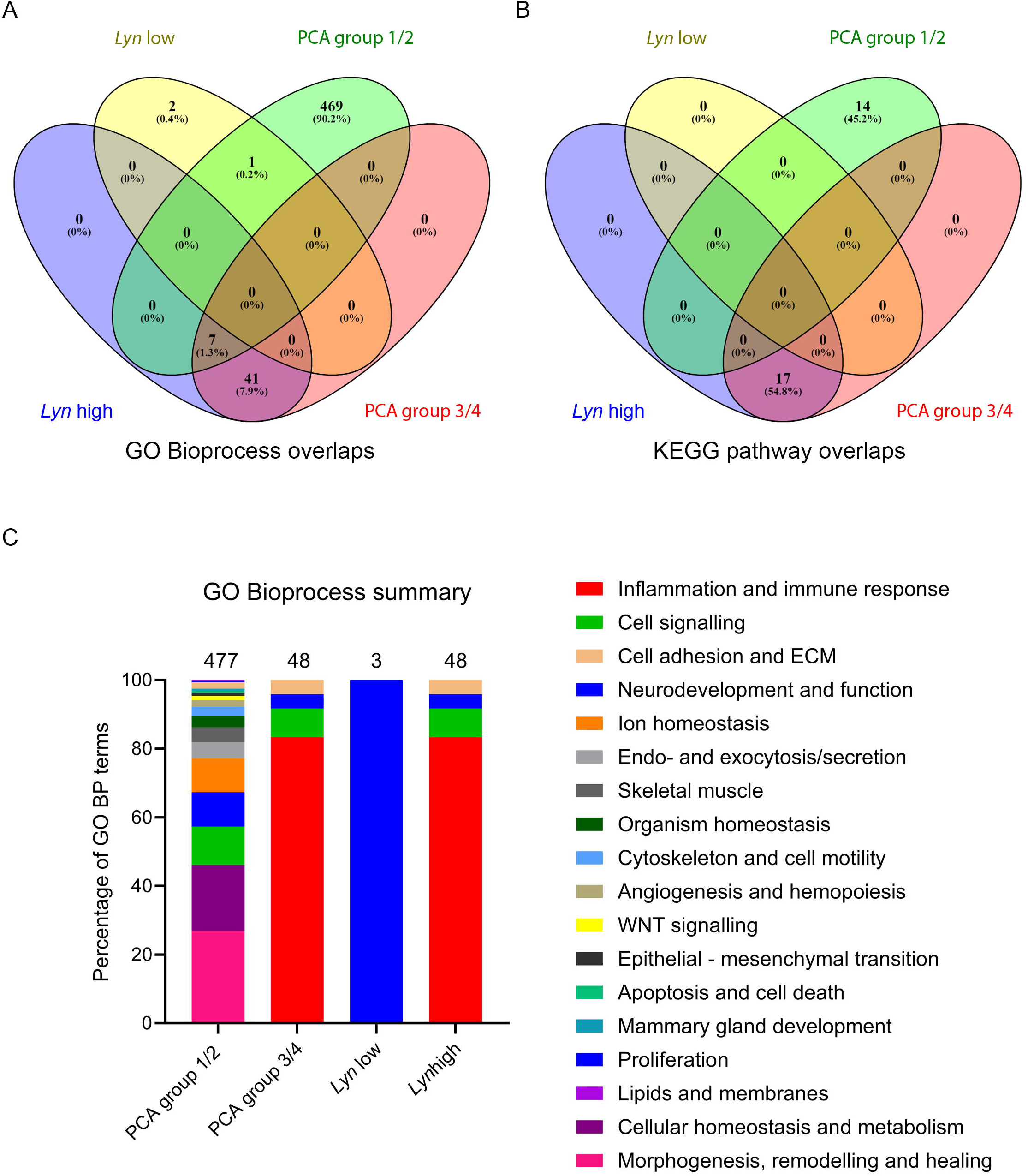
The *Lyn* high and PCA group 3/4 tumours are enriched for identical biological functions. **A)** Venn diagram analysis of overlap between GO Bioprocess terms enriched in the DEGs from *Lyn* high, *Lyn* low, PCA group 1/2 and PCA group 3/4 tumours. **B)** Venn diagram analysis of overlap between KEGG pathways enriched in the DEGs from *Lyn* high, *Lyn* low, PCA group 1/2 and PCA group 3/4 tumours. **C)** Distribution of enriched GO Bioprocess terms within functional categories for the *Lyn* high, *Lyn* low, PCA group 1/2 and PCA group 3/4 tumours. The number of enriched GO BP terms identified in the DEGs of each tumour group is indicated above each bar. Each bar is divided up according to the percentage of enriched GO BP terms falling into each functional category (indicated by the colour key).

Considering that the PCA group 3/4 tumours had elevated *Lyn*/LYN expression (**Fig. 5B,C**), we next assessed the overlap between the sets of significant DEGs from the *BlgCre Brca1^fl/fl^ p53^+/-^ Lyn^wt/wt^* vs. *BlgCre Brca1^fl/fl^ p53^+/-^ Lyn^fl/fl^* comparison, the *Lyn* high vs. *Lyn* low tumour comparison and the PCA group 3/4 vs. PCA group 1/2 comparison, using Venny (https://bioinfogp.cnb.csic.es/tools/venny/) (**Fig. 5F**). There was very little overlap between the *BlgCre Brca1^fl/fl^ p53^+/-^ Lyn^wt/wt^* vs. *BlgCre Brca1^fl/fl^ p53^+/-^ Lyn^fl/fl^* DEGs and the other comparisons. However, 340 genes were identified as significantly differentially expressed in both the *Lyn* high vs. *Lyn* low and the PCA group 3/4 vs. PCA group 1/2 comparison. Of these, 282 were upregulated in both *Lyn* high and PCA group 3/4 tumours while 57 were upregulated in both *Lyn* low and PCA group 1/2 tumours. Only one gene was elevated in *Lyn* high and PCA group 1/2 tumours (**Table S10**).

The distribution of the *Lyn* high and *Lyn* low tumours within the PCA groups was consistent with these results. Of the PCA group 3/4 tumours, six were also in the *Lyn* high tumour group while four were tumours with intermediate *Lyn* expression. There were no *Lyn* low tumours in PCA group 3/4. PCA group 1/2 included all the *Lyn* low tumours, nine intermediate *Lyn* expression tumours and seven belonging to the *Lyn* high group. Notably, the *Lyn* high tumours in PCA group 3/4 were six of the seven tumours with the highest rank for *Lyn* expression by RNAseq, the one exception being one of the *Lyn* high tumours in PCA group 1/2 which ranked second in *Lyn* expression by RNAseq but had been scored 0 for LYN expression by IHC (**Table S5**).

### *Lyn* high and PCA group 3/4 tumours are enriched in inflammatory signalling pathways while PCA group 1/2 tumours are enriched in morphogenesis and cancer-associated signalling pathways

We next undertook Gene Set Enrichment Annotation (GSEA) of the DEGs from the *Lyn* high versus *Lyn* low and the PCA group comparisons using g:Profiler. Genes significantly differentially expressed in the two comparisons were annotated separately and then overlaps between the annotations assessed. Full details are provided in **Table S10**. For ease of interpretation, we concentrated on understanding differentially enriched Gene Ontology Bioprocess terms (GO BP terms) and KEGG pathways. GO BP terms were grouped by functional categories to facilitate this. The GO BP and KEGG analysis is summarised in **Table S11** and **Fig. 6**.

*Lyn* high tumours were enriched for 48 GO BP terms and 17 KEGG pathways. *Lyn* low tumours were enriched for 3 GO BP terms but no KEGG pathways. PCA group 1/2 tumours were enriched for 477 GO BP terms and 14 KEGG pathways while PCA group 3/4 tumours were enriched for 48 terms and 17 KEGG pathways (**Table S11**). The overlaps in GO BP and KEGG pathways between the tumour groups was striking and reflected the overlap seen in the DEGs. The list of enriched GO BP and KEGG pathways in the *Lyn* high and PCA group 3/4 tumours was identical. Seven of these GO Bioprocesses were also enriched in PCA group 1/2, however, the majority (469 out of 477) of GO BP terms and all KEGG terms enriched in PCA group 1/2 were not found in the other groups (**Fig. 6A, B**).

GO BP terms were categorised into functional classes (**Table S11**) and the proportions of enriched terms from each functional class assessed for the tumour groups (**Fig. 6C**). Unsurprisingly, given that the terms enriched in the *Lyn* high and PCA group 3/4 tumours was identical, the classification of GO BP terms in these groups was identical. For both of these sets, the most numerous classification of enriched GO BP terms was inflammation and immune response (40 terms, 83.3%), followed by cell signalling (4 terms, 8.3%), cell adhesion and ECM (2 terms, 4.2%) and neurodevelopment and function (2 terms, 4.2%). For the *Lyn* low tumours, the three enriched GO BP terms were all classified as associated with neurodevelopment and function (100%). The 477 GO BP terms enriched in PCA group 1/2 could be classified into 17 different functional classes. The four largest of these (to which >10% of the 477 GO BP terms were assigned) were morphogenesis, remodelling and healing (128 terms, 26.8%), cellular homeostasis and metabolism (92 terms, 19.3%), cell signalling (53 terms, 11.1%) and neurodevelopment and function (48 terms, 10.1%).

We next examined the enriched KEGG pathways. Consistent with the GO BP analysis, and the overlap of the annotations between the tumour groups, the *Lyn* high/PCA group 3/4 tumours were enriched for KEGG pathways including ‘Cytokine-cytokine receptor interaction’, ‘NF-kappa B signaling’, ‘Chemokine signaling’, ‘TNF signaling pathway’ and ‘Apoptosis’. In contrast, the PCA group 1/2 tumours were enriched for KEGG pathways including ‘Breast cancer’, ‘Wnt signaling’, ‘Notch signaling’, ‘PI3K-Akt signaling’ and ‘Pathways in cancer’ (**Table S11**).

### *In vivo* differences between tumour groups are not maintained in cultured neoplastic cells

We next carried out qrtPCR analysis of expression of seven inflammation and immunity/NfkB-associated genes (*Bcl2a1a*, *Cd40*, *Nfkb2*, *Relb*, *Ccl5*, *Tnfaip3*, *Shisa8*) differentially expressed between the *Lyn* groups and PCA groups of tumours, using twenty of the samples (**Table S12**) analysed by RNAseq, in order to validate the analysis. The results confirmed that the genes were significantly differentially expressed between the *Lyn* low and *Lyn* high (**Fig. 7A; Table S12; Fig. S7**), and the PCA 1/2 and PCA 3/4 tumour groups (**Fig. 7B**) and in expected patterns (*Bcl2a1a*, *Cd40*, *Nfkb2*, *Relb*, *Il4i1*, *Ccl5* and *Tnfaip3* significantly more highly expressed in *Lyn* high and PCA 3/4 tumours, *Shisa8* significantly more highly expressed in *Lyn* low and PCA 1/2 tumours).

**Figure 7:**
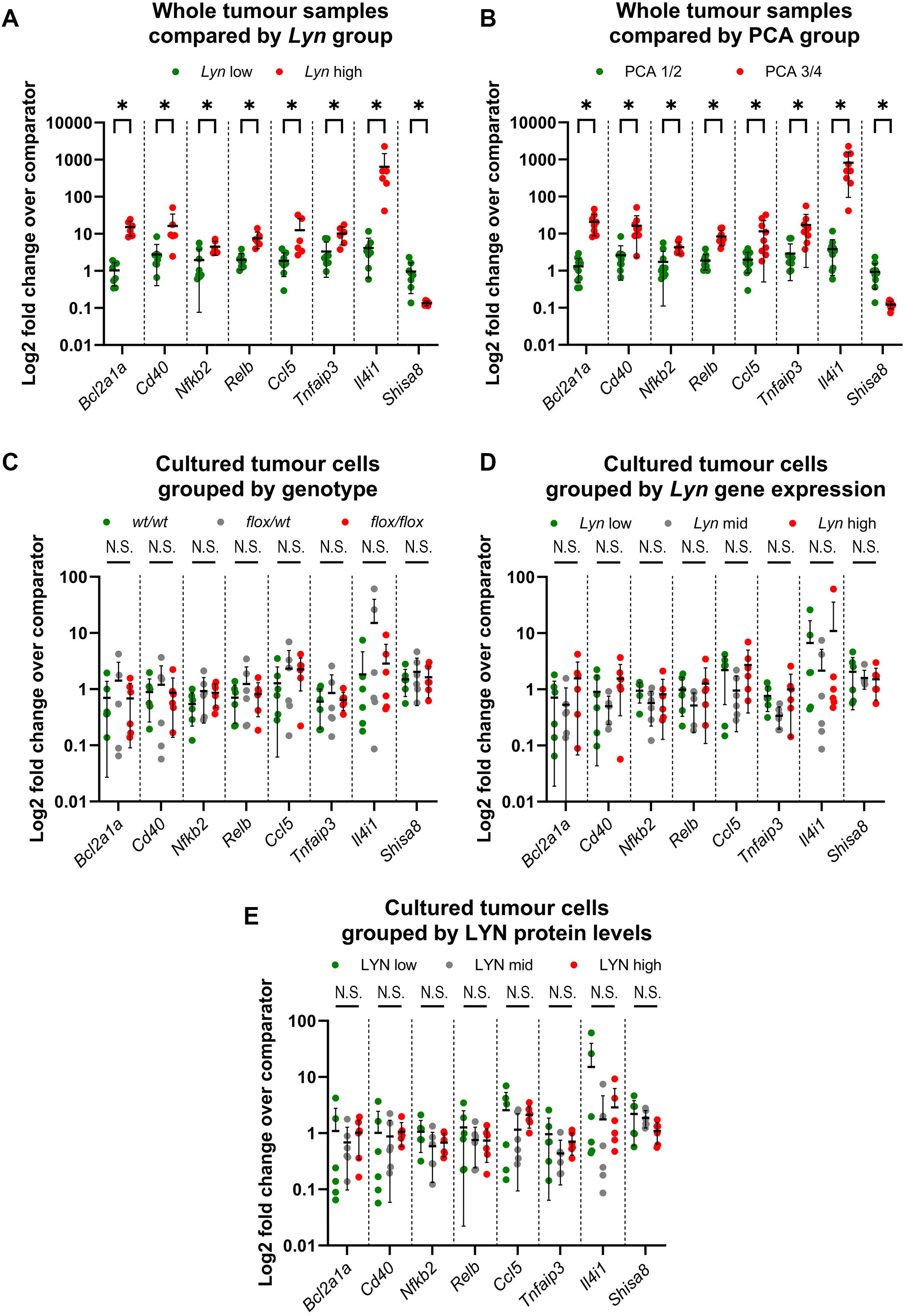
Gene expression differences between tumour in vivo are not maintained in neoplastic tumour cells in primary culture. **A,B)** Quantitative real-time rtPCR (qrtPCR) validation of *Ccl5*, *Tnfaip3*, *Bcl2a1a*, *Cd40*, *Nfkb2*, *Relb*, *Shisa8* and *Il4i1* expression in whole tumour samples analysed by RNAseq, comparing *Lyn* low (n=8) and *Lyn* high (n=6) **(A)** and PCA 1/2 (n=11) and PCA 3/4 (n=9) **(B)** tumour groups. Patterns of gene expression are consistent with the RNAseq data. **C-E)** qrtPCR expression of the same gene set in primary cultures of neoplastic tumour cells from *Lyn^wt/wt^*, *Lyn^flox/wt^* and *Lyn^flox/flox^* tumours (n=6 of each genotype). Expression is compared by genotype **(C)**, previously determined levels of *Lyn* gene expression **(D)** or previously determined levels of LYN protein expression **(E)** (Figure 3). Cultures divided into three groups based on high (top third), mid (middle third) or low (bottom third) levels of expression. There are no differences between any groups in the cultured cell analysis. Data presented as expression levels normalised to *Gapdh* and *B-actin* (A,B) or *Gapdh* alone (C-D) (**Figure S6**) and relative to comparator samples (**Supplementary Table 12**). Mean±s.d., Mann-Whitney tests with multiple comparison correction, *P<0.05, N.S., not significant.

Next, we analysed expression of the same set of genes in the set of tumour cell primary cultures previously analysed for *Lyn* conditional allele recombination and *Lyn* gene and LYN protein expression (**Fig. 3**). In contrast to the results from the whole tumour analysis, there were no significant differences between cultured cells, whether one compared cultures from different cohorts (**Fig. 7C**), cultures with different levels of *Lyn* gene expression (**Fig. 7D**) or cultures with different levels of LYN protein expression (**Fig. 7E**).

### B-cell abundance in tumours correlates with doubling time

The results of the qrtPCR validation suggest that either the differences in tumour biology suggested by the RNAseq analysis are not a result of differences between the neoplastic cells in the tumours but are a consequence of other cell types, or differences only appear between the neoplastic cells when they are in the context of an *in vivo* microenvironment. A combination of the two factors is also possible.

One potential difference between the *Lyn* low/PCA group 1/2 and *Lyn* high/PCA group 3/4 tumours is the extent of immune cell infiltration. This possibility is supported by immunoglobulin genes being significantly more highly expressed in the *Lyn* high/PCA group 3/4 tumours (**Table S9**), the enrichment of this tumour group for ‘inflammation and immunity’ associated genes (**Fig. 6**) and the known high expression of *Lyn* in immune cell subsets, particularly B-cells (Brian and Freedman, 2021). Therefore, to assess the immune cell infiltration between the tumour groups we analysed the RNAseq data using CIBERSORTx (Steen et al., 2020).

There were no significant differences in immune cell subsets between the tumour groups (**Fig. S7; Table S13**). However, when considering memory B-cells and plasma cells in particular (two subsets likely to contribute significantly to a ‘high *Lyn*’ ‘high *Ig* gene’ signature), the *Lyn* high / PCA group 3/4 tumours had a higher mean abundance of cells than the *Lyn* low / PCA 1/2 group, but also with large error bars (*Lyn* low tumours memory B-cells abundance 30.007±35.411, mean±s.d., n=13; *Lyn* high tumours memory B-cells abundance 126.619±119.442, mean±s.d., n=13; *Lyn* low tumours plasma cells abundance 1.614±2.689, mean±s.d., n=13; *Lyn* high tumours plasma cells abundance 41.141±51.980, mean±s.d., n=13; t-tests fail to meet significance threshold following correction for multiple testing across the CIBERSORTx dataset) (**Fig. S7; Table S13**). Therefore, while a subset of *Lyn* high / PCA group 3/4 tumours did have high levels of immune cells likely to contribute to tumour gene expression signatures, this was not true of all of them.

*Lyn* high / PCA 3/4 tumours had a higher histoscore than *Lyn* low / PCA 1/2 tumours (**Figs. 4D and 5C**), so we next assessed whether there was an association between LYN staining of tumour cells, as assessed by histoscore, and B-cell abundance. The group of tumours with the highest histoscore included four tumours with the highest abundance of B-cells. However, the high LYN tumour cell histoscore tumours also included tumours with low or no B-cell infiltrate and there was no significant difference in abundance overall between tumours with different levels of LYN staining (**Fig. 8A,B**).

**Figure 8:**
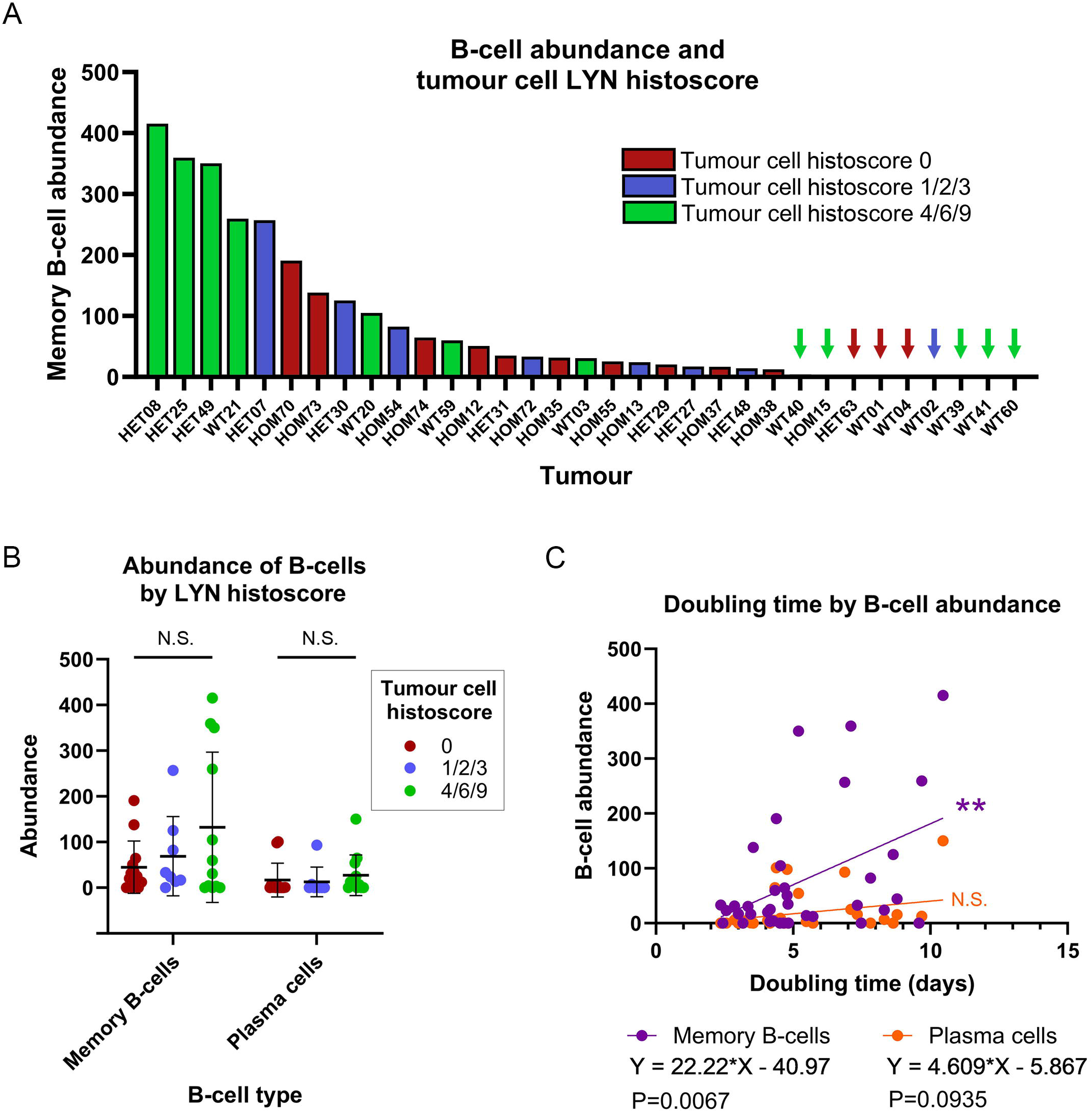
Memory B-cell abundance correlates with tumour doubling time. **(A)** Memory B-cell abundance in 33 RNAseq tumour samples plotted from highest to lowest abundance (arbitrary units) and colour-coded by tumour cell LYN histoscore for the tumour. Very low/zero abundance samples are indicated by colour-coded arrows. **(B)** Memory B-cell and plasma cell abundance compared by tumour cell LYN histoscore groups (n=13, 8, 12, for groups 0, 1/2/3, 4/6/9 respectively). Mean±s.d.. No significant differences (ANOVA). **(C)** Simple linear regression of B-cell abundance (arbitrary units) (Memory B-cell, purple; Plasma cells, orange) against tumour double time (days) (n=36). Increased numbers of Memory B-cells are significantly associated with increased tumour doubling time.

Finally, we determined whether there was an association between B-cell abundance and tumour doubling time. Indeed, there was a significant association between memory B-cell (P=0.0067) but not plasma cell (P=0.0935) abundance and tumour doubling times (**Fig. 8C**).

## Discussion

The SRC-family kinase LYN is most highly expressed in haematopoietic cells but is also expressed in a wide variety of other tissues, including epithelia (reviewed in (Brian and Freedman, 2021)). It is a downstream target of c-KIT signalling in luminal epithelial stem/progenitor cells of the mammary gland (Regan et al., 2012; Tornillo et al., 2018) and also expressed in breast cancers, particularly TNBC (Choi et al., 2010; Hochgrafe et al., 2010; Molyneux et al., 2010).

LYN kinase is best known for its role as both a positive and negative regulator of myeloid and B-cell development and differentiation (Brian and Freedman, 2021). The positive functions of LYN are context dependent and in positive signalling loss of LYN may be compensated for by other SRC-family kinases. In contrast, LYN appears to be absolutely required for negative regulation of B-cell proliferation (Brian and Freedman, 2021; Xu et al., 2005). Notably, both *Lyn* knockout mice and LYN constitutively over-expressing mice develop lethal autoimmune kidney disease, although of distinct pathologies (Hibbs et al., 2002; Hibbs et al., 1995).

LYN has two splice isoforms (LYN^FL^/LYN p56/LYNA and LYN^Δ25-45^/LYN p53/LYNB) (Brian and Freedman, 2021; Tornillo et al., 2018; Xu et al., 2005) and a ΔNLYN caspase-cleaved variant which affects NFkB signalling has also been described (Marchetti et al., 2009). These splice isoforms are still not fully understood, although it is known that LYNA regulates a signalling checkpoint in macrophages (Freedman et al., 2015) and re-expression of either LYNA or LYNB in *Lyn* knockout mice restores B-cell developmental defects but neither rescues the autoimmune phenotype on its own (Brian et al., 2022).

Previous studies on the mammary epithelium and breast cancer, including our own, have highlighted LYN as a positive regulator of cell growth and survival (Choi et al., 2010; Tornillo et al., 2018). We have shown LYNA specifically regulates cell invasion and migration in TNBC cell lines *in vitro* but both LYNA and LYNB enhance breast cancer cell line survival, suggesting it is an oncogene (Tornillo et al., 2018). The tyrosine residue (Y32) in the LYNA-specific N-terminal region is a target of EGFR kinase activity and, once phosphorylated, results in LYN-mediated activation of the MCM7 DNA replication licencing factor (Huang et al., 2013). LYN activity has been reported to promote EMT through Vav-Rac1-PAK1-mediated control of SNAI protein localisation and stability in multiple cancer types, including breast cancer (Thaper et al., 2017). A role for LYN in regulation of p53 has also been described. LYN is reported to directly interact with p53 and prevent its nuclear export, suppressing MDM2-mediated p53 degradation and enhancing p53-dependent apoptosis (Ren et al., 2002).

Here, we find contradictory evidence for the role of LYN in mammary tumours, potentially related to functions in multiple cell types within a tumour. LYN protein expression was decreased in epithelial-origin neoplastic tumour cells carrying two copies of a conditional knockout *Lyn* allele in which CRE recombinase expression was under the control of the *Blg* promoter (**Fig. 3C**). However, some tumours retained strong LYN expression, suggesting incomplete recombination *in vivo*; furthermore, some tumours with wild type alleles showed very low levels of expression. Importantly, there was no association between LYN protein expression (by histoscore) and tumour doubling time (**Fig. 4F**).

In contrast, there was an association between *Lyn* expression levels as measure by RNAseq in whole tumour lysates and tumour doubling time (tumours with higher overall *Lyn* expression grew more slowly; **Fig. 4E**) and when tumours were grouped in an unsupervised manner on the basis of RNAseq data, there was also a correlation with doubling time (**Fig. 5D**). Therefore, overall *Lyn* expression correlated with tumour doubling time, but LYN expression specifically in the tumour cells did not. Rather, tumour doubling time correlated with B-cell abundance (as defined by CIBERSORTx, which has been previously proven to be robust) (Steen et al., 2020) with tumours having a higher B-cell abundance score growing more slowly (**Fig. 8C**); there tended to be more B-cells in *Lyn* high / PCA group 3/4 tumours, although with considerable variation (**Fig. S7**). As B-cells are known to express *Lyn*, their presence in a tumour would tend to result in tumours with a heavy B-cell infiltration being grouped in the *Lyn* high / PCA 3/4 set (and in this tumour set having significantly higher levels of expression of inflammation and immunity genes).

As there was an association between tumour cell LYN histoscore and the *Lyn* tumour group by RNAseq (**Fig. 4D**, **Fig. 5D**), high *Lyn* expression by RNAseq analysis of whole tumour lysates likely resulted from a combination of moderate or high levels of *Lyn* transcript in the neoplastic tumour cells themselves as well as varying degrees of B-cell infiltrate, in some cases in very high abundance. It was the B-cell infiltrate which correlated with the doubling time of the tumours, rather than levels of LYN expression in the tumour cells.

However, it may be that LYN-dependent signalling pathways in tumour cells activated intrinsic inflammatory signalling pathways, potentially including NFkB, resulting in production of cytokines which enhanced immune cell recruitment and an anti-tumour immune response. Downstream mediators linked to LYN activation of NFkB include MEK, IKKα (Cooper et al., 2013), PI3K (Toubiana et al., 2015), MAPK and IkB (Avila et al., 2012) and there is also support for a role of NFkB activation in BRCA1 loss-of-function-associated breast cancers. NFkB activation was proposed to be the mechanism underlying hormone-independent growth of *BRCA1*-deficient luminal progenitors in colony formation assays *in vitro* (Lim et al., 2009; Sau et al., 2016). In contrast, a subset of *BRCA1*-mutant breast cancers were reported to show increased NFkB activity correlating with good prognosis (Buckley et al., 2016). These tumours were associated with increased numbers of CD8+ cytotoxic T-cells, suggested to create an ‘anti-tumour microenvironment’ (Buckley et al., 2016). The difference between these *in vitro* and *in vivo* studies reflects our own findings.

Our manuscript has limitations. In particular, LYN staining may not reflect active LYN protein or active LYN-dependent signalling. Unfortunately, antibodies are not currently available for immunohistochemistry which are specific for active pLYN. Those that are available stain the phosphorylated active site of all SRC-family kinases. Flow cytometry to purify neoplastic tumours cells for analysis by, for example, western blot, is also problematic. Antibodies are not available which can be used to specifically mark all neoplastic epithelial-origin cells as opposed to non-neoplastic epithelium, or indeed other components of the tumour. In the absence of a robust approach to purifying tumour cells, we opted to carry out RNAseq analysis from pieces of tumour which likely contained mixed populations of cells. Such pieces were taken from tumour regions away from obvious necrosis, but without any other selection criteria. This has the advantage of ensuring that sensitive RNA expression patterns are not altered during cell purification protocols but given that *Lyn* expression in *Lyn* high tumours was a result of, as we now propose, both LYN-expressing tumour cells and immune cells, interpretation of the results is complex. Future studies to test our model that activity of LYN-dependent, cell intrinsic signalling pathways results in the recruitment of an anti-tumour immune response will likely require single cell transcriptomic analysis or a similar approach.

Overall, our study suggests that, despite previous evidence supporting a cell-intrinsic role for LYN kinase in promoting mammary tumour cell survival, proliferation and invasion, its potential role in B-cells and anti-tumour immune response means that as a therapeutic target in breast cancer LYN kinase is likely to present difficulties. We also suggest that previous studies reporting associations between *Lyn* overexpression in TNBC may actually be reflecting enrichment in TNBC for immune cells, as is being exploited by current immunotherapy trials in this setting (Tarantino et al., 2022).

## Materials and Methods

See **Table S1** for a full list of all primers, antibodies and other reagents. Raw scanned western blots are provided as a **Supplemental Data File**.

### Establishment of genetically modified mouse lines

This study was approved by the Cardiff University Animal Welfare and Ethical Review Body and carried out under the authority of appropriate Home Office Personal and Project Licences and with reference to ARRIVE guidelines (Percie du Sert et al., 2020). In particular, animals were monitored regularly and predefined humane endpoints strictly adhered to. Numbers of animals required in each cohort were based on previous experience of requirements for a sufficiently powered tumour cohort study (which are typically 15 – 20 animals per cohort depending on effect size, but given the inherent random nature of litter sizes, sex ratios and genotypes, numbers in each cohort may not be identical). Randomisation was not appropriate as animals had to be assigned to cohorts according to their genotype. Only female animals were used.

The full breeding scheme is illustrated in **Fig. S1**. Mice carrying the conditional *Brca1* allele on the *p53* heterozygote background as well as a CRE recombinase under the control of the β-lactoglobulin mammary specific promoter (*BlgCre Brca1^fl/fl^ p53^+/-^* mice) have been previously described (McCarthy et al., 2007; Molyneux et al., 2010). Mice carrying a tamoxifen-activated CRE ubiquitously expressed from the *Rosa26* locus (*R26C*) (Hameyer et al., 2007) were obtained from Prof. Karen Blyth, CRUK Beatson Centre, Glasgow. Mice carrying the *Lyn floxed exon 4* conditional allele (*Lyn^tm1c^*) were obtained from the Mary Lyon Centre, MRC Harwell (full nomenclature C57BL/6N-Lyn^tm1c(EUCOMM)Hmgu^/H, derived from an ES cell clone HEPD0704_6_B11).

The full details of all animals used in the study and of all histological samples are provided in Tables S2 and S3.

### Mouse mammary epithelial cell harvest and culture

Mammary epithelial organoids were prepared from fourth mammary fat pads of 10-12 week-old virgin female mice as described (Smalley, 2010). Intramammary lymph nodes were removed prior to tissue collection. Fat pads were finely minced on a McIlwain Tissue Chopper and then digested for 1 hr at 37°C in 3 mg/ml collagenase A/1.5 mg/ml trypsin in serum- and phenol red-free L15 medium (ThermoFisher Scientific / Invitrogen, Paisley, UK) with gentle rotation. Tissue fragments (‘organoids’) released were incubated for 5 min in Red Blood Cell Lysis buffer (Merck Sigma-Aldrich, Gillingham, Dorset, UK), washed and then plated for 1 hr at 37°C in DMEM/10%FBS (ThermoFisher) for depletion of fibroblasts by differential attachment.

For three-dimensional (3D) cultures, organoids were incubated with 0.05% trypsin/EDTA for 2 min at 37°C prior to plating onto Growth Factor Reduced phenol red-free Matrigel (Fisher Scientific, Loughborough, Leicestershire, UK) in complete growth medium (DMEM:F12 with 10% Charcoal Stripped FBS, 5 ug/ml insulin, 10 ng/ml cholera toxin and 10 ng/ml epidermal growth factor (Merck Sigma-Aldrich). 4-Hydroxytamoxifen (4OHT) (Merck Sigma-Aldrich) was added at a final concentration 100 nM for 10-12 hours to induce the recombination of the *Lyn^fl^*allele.

### Isolation of Primary Tumour Cells

Primary tumour epithelial cells were obtained using the gentle MACS Dissociator and Mouse Tumor dissociation kit (Miltenyi Biotec, Bisley, Surrey, UK) following the protocol recommended for ‘Dissociation of Tough Tumors’. To ensure efficient dissociation, volumes of Enzyme D, Enzyme R and Enzyme A were scaled up according to the size of the tumour piece (100 uL, 50 uL and 12.5 uL respectively per each 0.5 cm^3^). The optional red blood cell lysis step was included in the procedure. Resulting cells were plated in complete growth medium in two-dimensional (2D) adherent conditions. Cells at passage 0 were used for all the experiments in this study.

### Tumour measurements and doubling times

Tumour width (W) and length (L) were measured using a caliper twice a week by the same person each time to eliminate inter-operator variability. Volume was calculated using the formula (L x W^2^/2).

### IHC and FFPE sample processing

Mice were euthanised by an approved method when previously established humane endpoints were reached. A full necropsy was performed and any tumour tissue was fixed in 10% neutral buffered formalin for 24h at 4°C before being processed into paraffin blocks according to standard procedures. When a tumour was of sufficient size, a piece (distant from any obvious necrosis) was also snap frozen on dry ice at time of dissection and then stored at −80°C for later RNA/protein extraction. In some cases, pieces of tumour were kept in L15 medium on ice for later isolation and culture of primary tumour cells. Visceral organs (liver, kidneys, spleen, lungs and in some cases heart and stomach, if obvious pathology present) were also fixed in neutral buffered formalin for 24 h and processed into paraffin blocks.

Tissue sections (5 μm) were either stained using Haematoxylin & Eosin (H&E) for histological analysis or used for immunohistochemical staining. For the latter, freshly cut sections were dewaxed and re-hydrated. Sections underwent antigen retrieval in citrate buffer, pH 6.0 in a pressure cooker for 15 min before incubation with a 3% hydrogen peroxide solution for 20 min and then blocking in 10% goat serum/0.1% Tween-20/TBS for 1 hour. Incubation with primary antibodies **(Table S1)** was performed overnight at 4°C. Detection was carried out using the ImmPRESS kit (Vector Labs, Peterborough, UK). Sections were counterstained with haematoxylin and mounted. Images were acquired using a VS200 slide scanner (Olympus Keymed, Southend-on-Sea, Essex, UK) with a 20x objective and visualised using OlyVIA slide viewer software (Olympus).

### Histopathological analysis

Mammary tumour phenotyping was carried out by MJS (who has over ten years’ experience of using the four-histotype classification system for mouse mammary tumours) using our previously established criteria based primarily on morphology of H&E stained sections and immunohistochemical staining for ΔNp63 (Melchor et al., 2014; Molyneux et al., 2010; Ordonez et al., 2023; Ordonez et al., 2019; Ordonez et al., 2021). In brief, assessment of metaplasia (either spindle cell or squamous) and the extent of any ΔNp63 staining allows mouse mammary epithelial tumours to be classified as adenosquamous tumours (ASQC; extensive squamous metaplasia and abundant ΔNp63 staining), adenomyoepitheliomas (AME; abundant ΔNp63 staining in a distinct pseudo-basal pattern bordering ΔNp63-negative cells, but little or no metaplasia), metaplastic spindle cell carcinomas (MSCC; extensive or near total spindle cell metaplasia with infrequent nests of epithelial tumour cells; no ΔNp63 staining), adenocarcinomas of no special type (ACNST; little or no metaplasia and little ΔNp63 staining). Histology of other organs was reviewed by MJS with support and advice from SB.

### Scoring of Ki67 IHC

For Ki67 IHC quantification, images of five different regions (in one case, six regions) from each section were captured using the OlyVIA software at 10x magnification. Regions were chosen to include areas with the highest level of Ki67 staining for that section, so that the final score represented the highest potential for proliferation, and therefore the most aggressive behaviour, of that tumour. The percentage of positive tumour cells in each image was determined automatically using Cognition Master Professional Ki67 Quantifier (Medline Scientific Limited, Chalgrove, Oxfordshire, UK). Values returned by the program were ‘sense-checked’ against each image; any obvious errors (e.g. 21-34-03 Field 5; **Table S4**) excluded from further analysis.

### Scoring of LYN IHC staining by modified histoscore

LYN IHC staining using a rabbit polyclonal antibody (Thermofisher) was quantified by a modified histoscore approach considering strength of staining and area stained. If staining was visible at x0.2 magnification on the OlyVIA slide viewer images, staining was scored as strength ‘3’. If staining was not visible at x0.2 but was visible at x2, it was scored as strength ‘2’. If staining was not visible at x2 but was visible at x20, it was scored as strength ‘1’. If no staining was visible at x20 it was scored as ‘0’. For the area of tumour stained, scoring was determined as follows: 0, no staining; 1, <10% of tumour cells positive; 2, 10 – 50% of tumour cells positive; 3, >50% of tumour cells positive. These divisions were chosen as easily assessable by eye, without the need for exact counting. The two scores were then multiplied together to give a final value. Note that the antibody used measures total LYN protein, not active protein. Antibodies specific to the phosphorylation site on LYN which indicates activation are not currently available.

### Protein isolation and western blot analysis

2D cultured cells were lysed in Laemmli buffer. 3D cultured primary mouse mammary cells were released from Matrigel using the BD cell recovery solution prior to lysis. Protein extracts were separated by SDS-PAGE on 4–15% gradient Mini-PROTEAN TGX Precast Protein Gels (Bio-Rad, Watford, Hertfordshire, UK), transferred to PVDF membranes (IPVH00010, Merck Millipore, Hertfordshire, UK) and immunoblotted with anti-LYN antibodies. GAPDH was used as loading control. Resulting immunocomplexes were detected by HRP-conjugated anti-mouse IgG or anti-rabbit IgG secondary antibodies and enhanced chemiluminescent (ECL) reagents (WBLUF0100, Merck Millipore).

### RNA isolation and gene expression analysis

Total RNA was isolated from tumour tissue or 2D-cultured cells using the RNeasy Minikit (QIAGEN) according to the manufacturer’s protocol. Trizol (ThermoFisher Scientific) was used for RNA extraction from 3D-cultured primary mammary organoids. Up to 1 ug of RNA was converted into cDNA using either the Quantitect Reverse Transcription kit (QIAGEN) or the Superscript IV transcription kit (ThermoFisher Scientific) following the manufacturer’s instructions. Gene expression analysis was carried out using either TaqMan Master Mix and Taqman gene expression assays (ThermoFisher Scientific) or Applied Biosystems SYBR Green Master Mix (ThermoFisher Scientific) and primers designed using Primer3 V4.1.0 (**Table S1**). Data analysis was carried out using the QuantStudio7 Software. Relative expression levels of target genes were calculated using the ΔΔCt method as described previously (Kendrick et al., 2008).

For validation of RNAseq analysis of whole tumours, the geometric mean of *Gapdh* and *B-actin* Ct values was used as a reference (Vandesompele et al., 2002). However, for analysis of passage 0 tumour cells in culture, only *Gapdh* was used as the reference, as analysis of *B-actin* variance in these cells suggested a batch effect which may have confounded the results (**Fig. S6**).

### RNA Sequencing and Analysis

Samples for RNAseq analysis underwent an on-column DNase I digestion step for genomic DNA removal prior to further processing. Total RNA quality and quantity was assessed using Agilent 4200 TapeStation and hsRNA or RNA ScreenTapes (Agilent Technologies, Stockport, Cheshire, UK). mRNA was isolated from 50ng of total RNA (RIN value >7) using the NEBNext® Poly(A) mRNA magnetic isolation module, (New England BioLabs, Hitichin, Herts, UK) (NEB, #E7490) and the sequencing libraries were prepared using the NEB® Ultra™ II Directional RNA Library Prep Kit for Illumina® (NEB, #E7760). The sequencing libraries were prepared following Chapter 1 of the NEB® Ultra™ II Directional RNA Library Prep Kit for Illumina® (New England BioLabs, NEB) protocol. The steps included mRNA isolation, fragmentation and priming, first strand cDNA synthesis, second strand cDNA synthesis, adenylation of 3’ ends, adapter ligation (1:80 dilution) and PCR amplification (14-cycles). Libraries were validated using the Agilent 4200 TapeStation and hsD1000 ScreenTapes (Agilent Technologies) to ascertain the insert size, and the Qubit® (Thermo Fisher Scientific) was used to perform fluorometric quantitation. The manufacturer’s instructions were followed except for the replacement of SPRIselect Beads or NEBNext Sample Purification Beads by AMPure XP beads (Beckman Coulter, High Wycombe, Herts, UK) in purification steps. The validated libraries were normalized to 4nM, pooled together and the pool sequenced on an S1 (200 cycle) flow cell using a 2×100bp PE dual index format on the Illumina® NovaSeq6000 sequencing system according to the manufacturer’s instructions. Samples were sequenced to a read depth of at least 35 Million prior to quality trimming with fastp (Chen et al., 2018). Quality trimmed reads were mapped to GRCm38 using STAR (v2.5.1b) (Dobin and Gingeras, 2015) with read multimapping filter set to 1 and gtf Gencode GRCm38 vM17. Exon and gene counts were calculated with featureCounts (v1.5.1) (Liao et al., 2014). Differential gene expression was calculated using SARtools using the DESeq2 package (Varet et al., 2016).

### Gene Set Enrichment Analysis (GSEA)

For GSEA, significantly differentially expressed genes from the different tumour groups (adjusted p value <0.05; log2 fold change <0.5 or >2.0) were uploaded to g:Profiler (https://biit.cs.ut.ee/gprofiler/gost) and queried against the *Mus musculus* database with a ‘term size limit’ of 1000 but otherwise using default options (with the exception of one group, the differentially expressed genes overlapping between the ‘PCA 1/2’ and ‘*Lyn* low’ tumour groups, for which the ‘term size limit’ was allowed default values, otherwise no results were returned). Results were downloaded as a CSV file (default options). Gene Ontology Bioprocess grouping into functional categories was carried out manually.

CIBERSORTx analysis (https://cibersortx.stanford.edu/) (Newman et al., 2019) was carried out using the LM22 signature matrix file for 22 immune cell types and was run in ‘absolute mode’ so that relative differences between the proportions of immune cell types would be maintained.

### Statistics

All statistical analysis was carried out in Prism 10.0.2 (GraphPad Software LLC). Survival curves were analysed by Log Rank test. For all other experiments, normally distributed data were analysed by ANOVA and/or t-tests where appropriate. Non-parametric data were analysed by Kruskel-Wallis and/or Mann-Whitney where appropriate. A P value of <0.05 was taken as significant. For analysis of difference in distribution of categorical variables, Chi2 test (two groups) or Chi2 test for trend (more than two groups) was used. P-values from multiple testing were corrected using the Holm-Sidak method.

Number of tumours available for analysis varied depending on the assay, as for some tumours doubling data were not available (as a minimum of three measurements was needed to determine this), for others IHC analysis was not available due to e.g. technical failures or poor quality / quantity of embedded material. For the RNAseq analysis, group sizes for determining differentially expressed genes varied depending on whether analyses were supervised or unsupervised. N numbers are provided in Figure legends.

## Acknowledgements

We acknowledge our colleagues at Wales Gene Park for their insight and expertise that assisted this research, for their technical and bioinformatic support in generating and analysing the NGS data. Wales Gene Park is an infrastructure support group with funding from Welsh Government through Health and Care Research Wales.

The authors would like to thank the Freedman Lab and also Catherine Hogan for their comments on the manuscript.

## Data availability

RNAseq data have been deposited at Gene Expression Omnibus (GEO) with the accession number GSE222075.

## Author contributions

Conceptualisation MJS, GT; Investigation GT, LW, HK, ATH, TH, MJS; Formal analysis GT, LW, HK, ATH, TH, SB, MJS; Writing – Original Draft, MJS, GT; Supervision, MJS; Project Administration, MJS; Funding Acquisition, MJS, GT.

