## Supplementary figures and images for "Conditional *in vivo* deletion of LYN kinase has little effect on a *BRCA1* loss-of-function-associated mammary tumour model"

### Figure S1

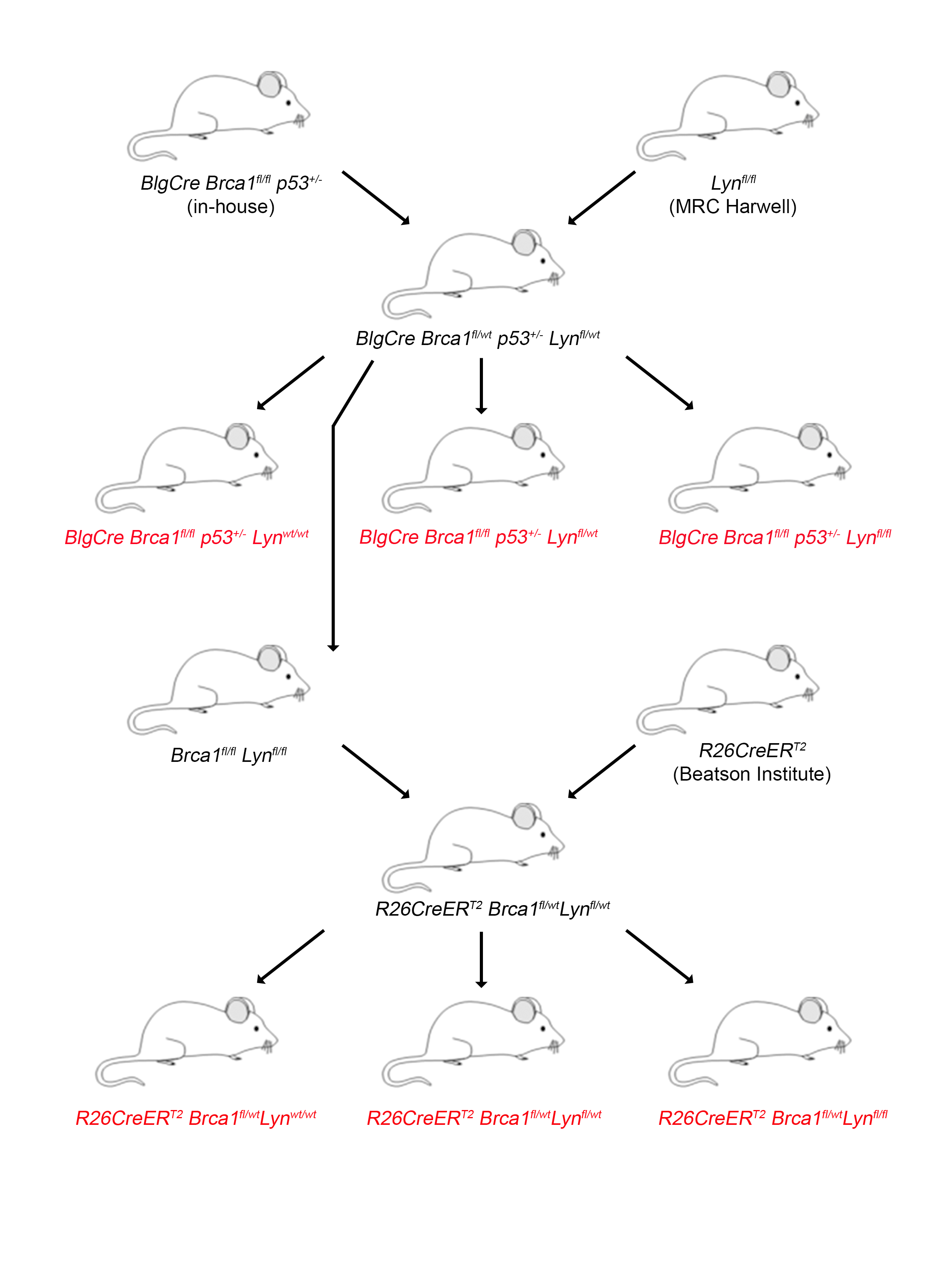

### Figure S2

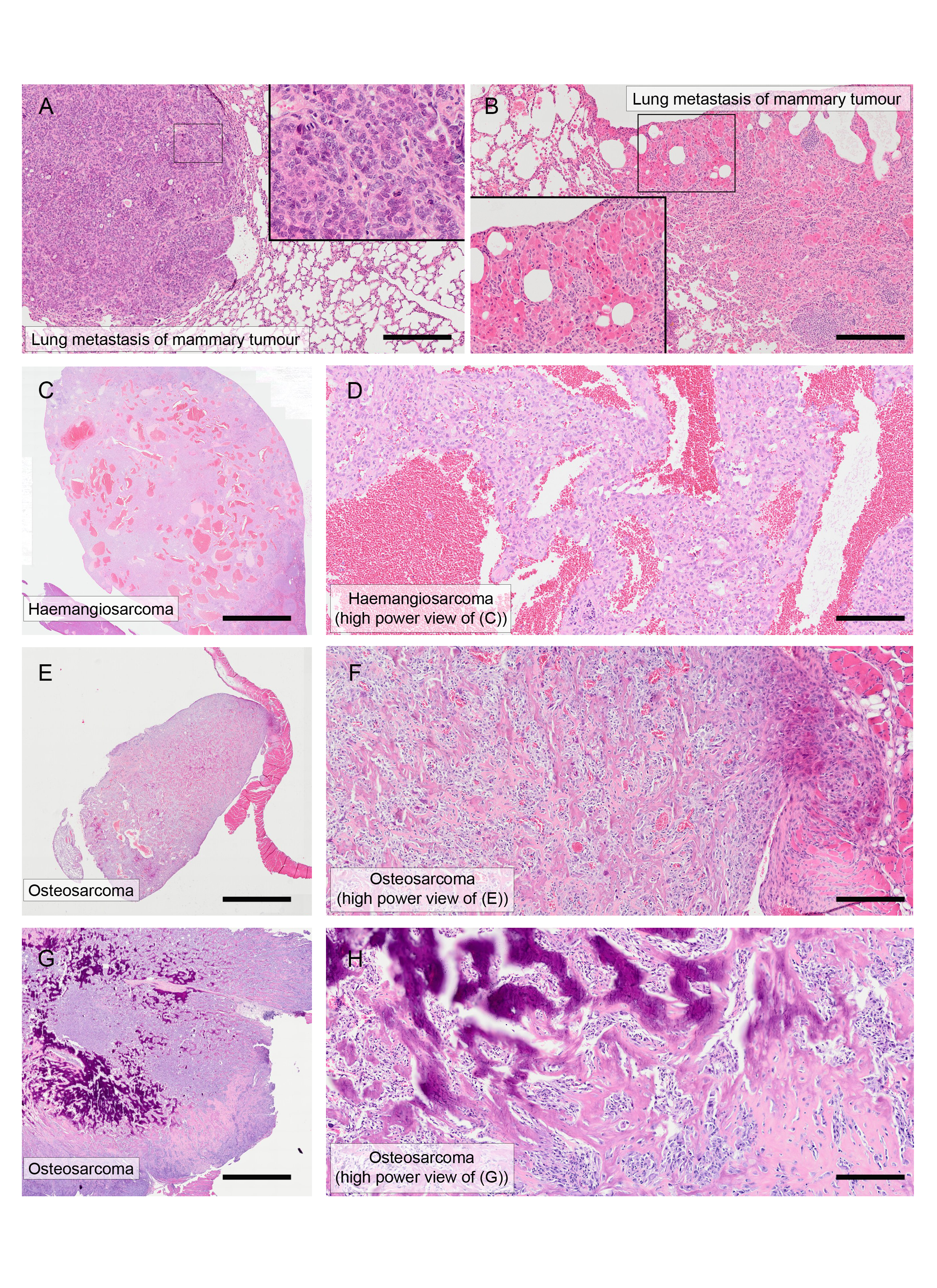

### Figure S3

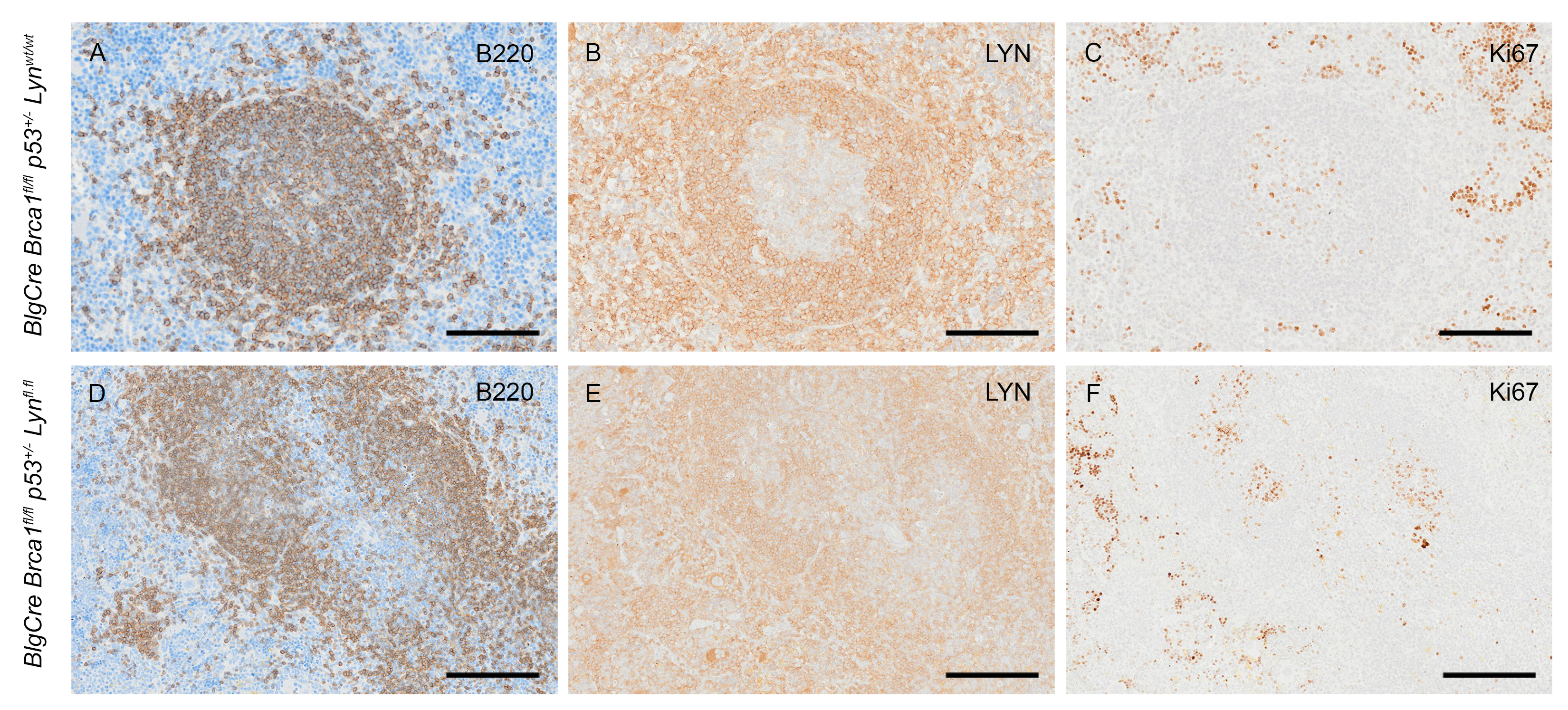

### Figure S4

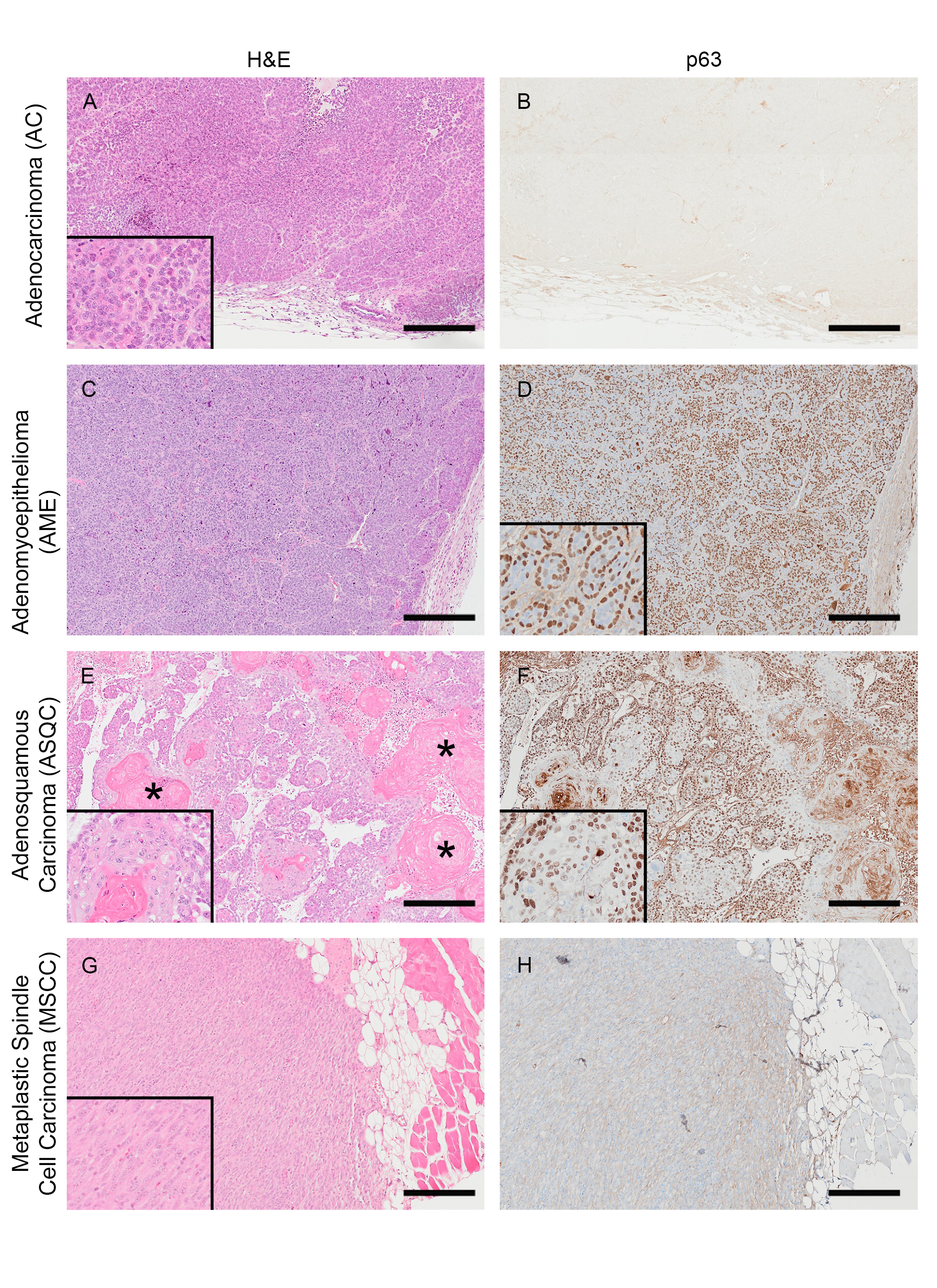

### Figure S5

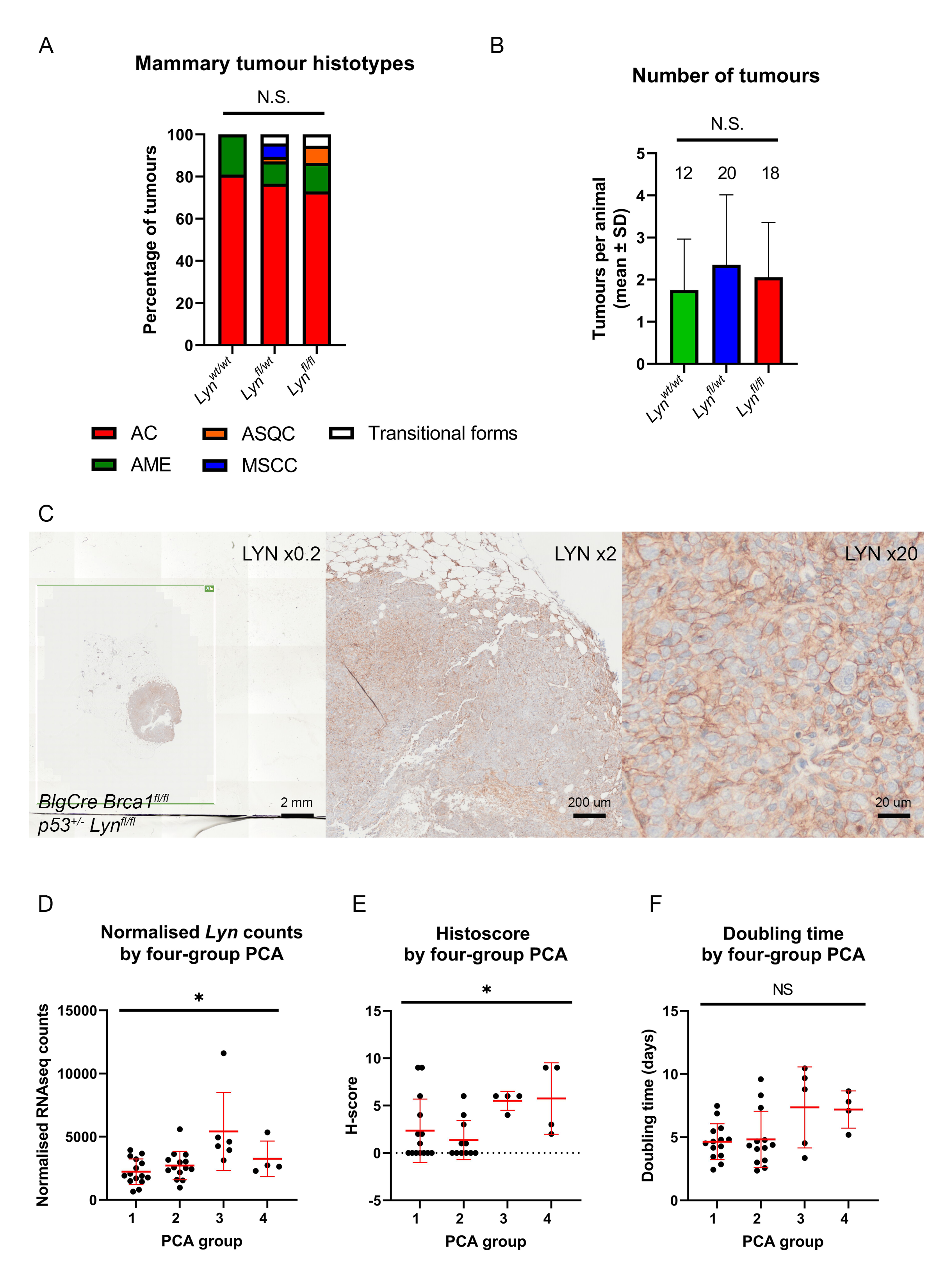

### Figure S6

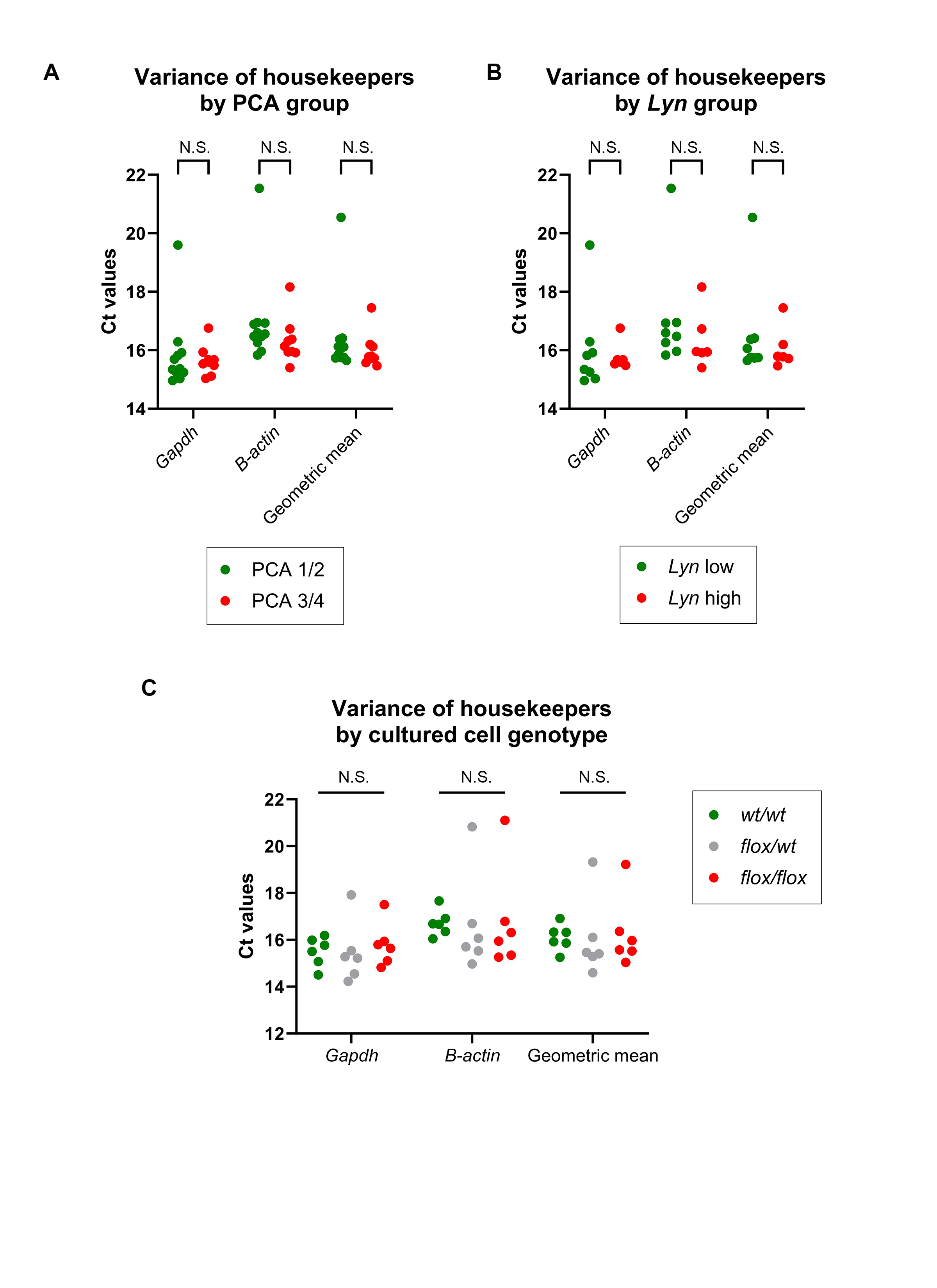

### Figure S7

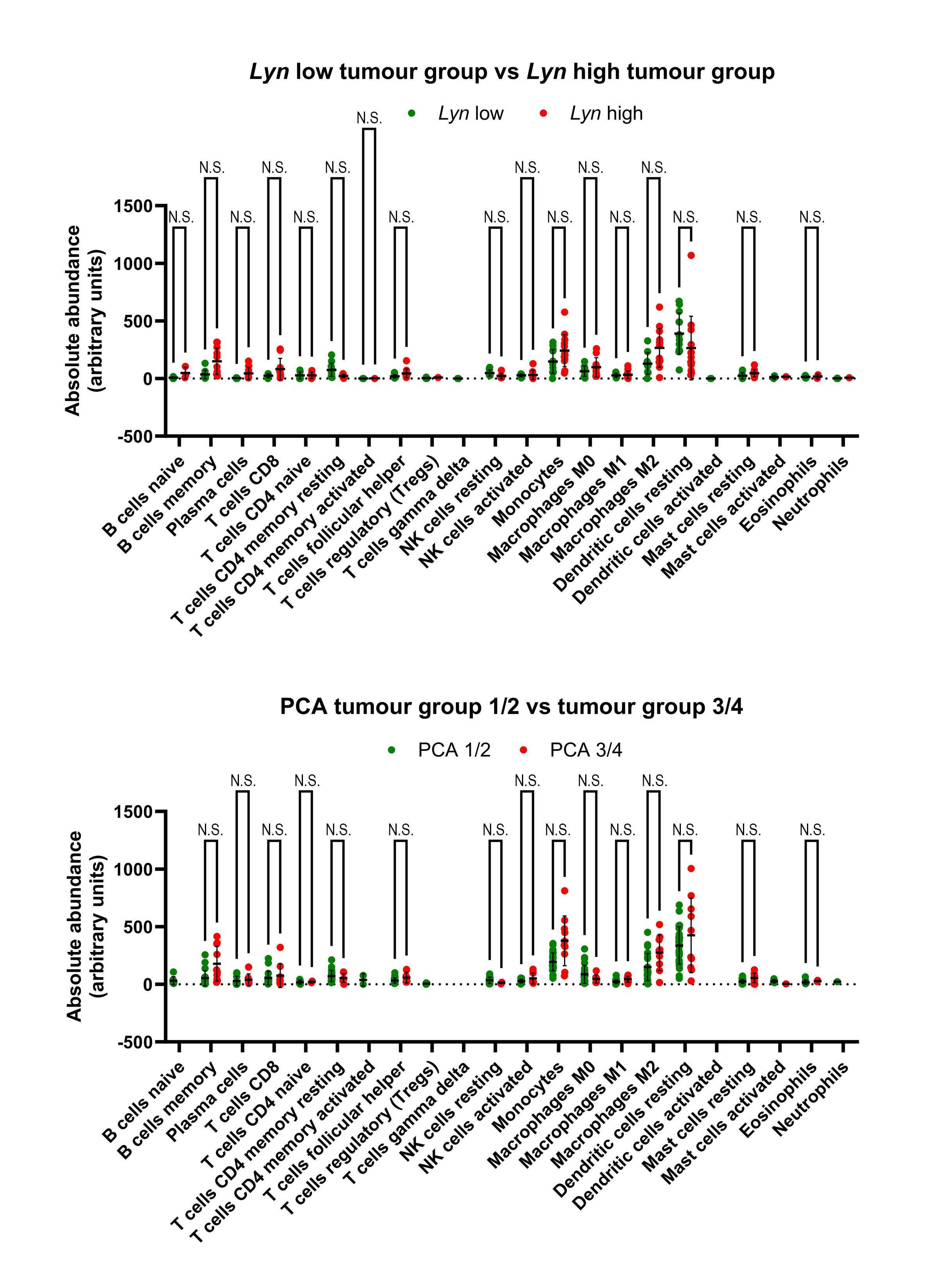

### Supplementary Data File

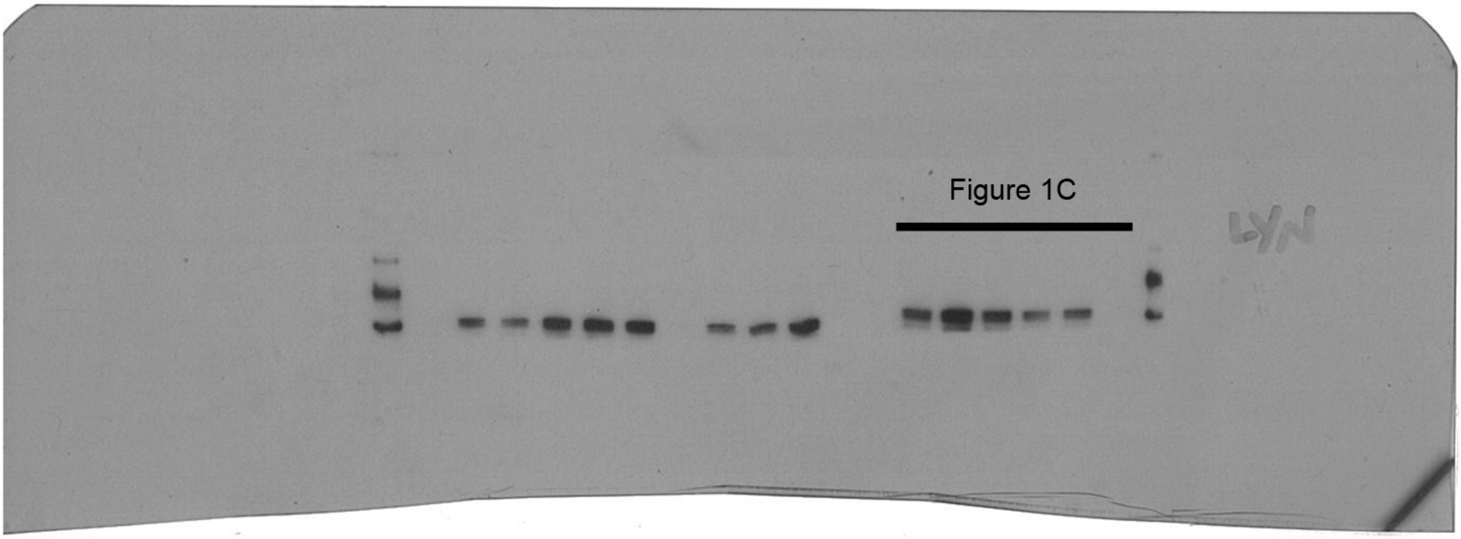

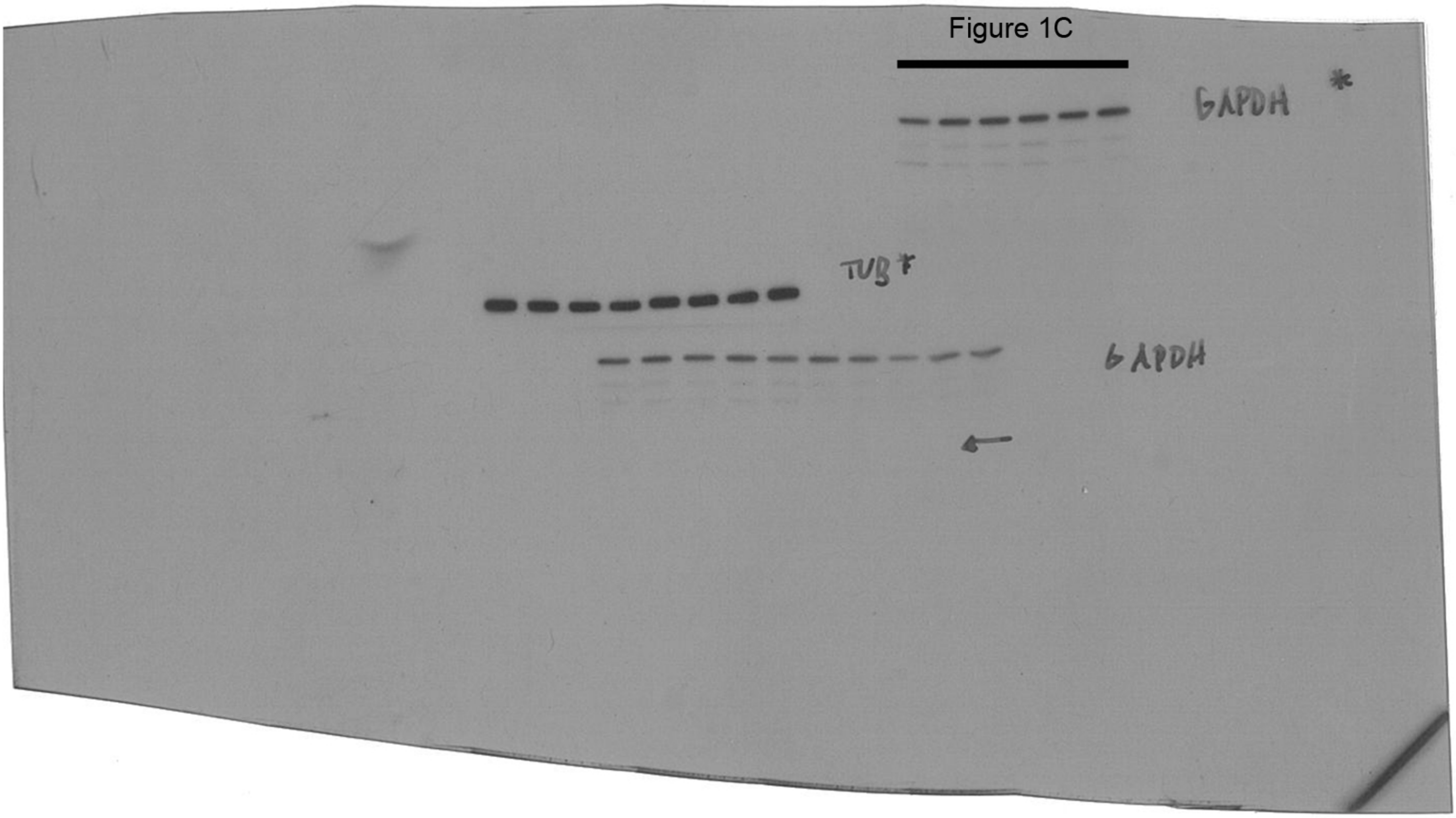

Figure 1D

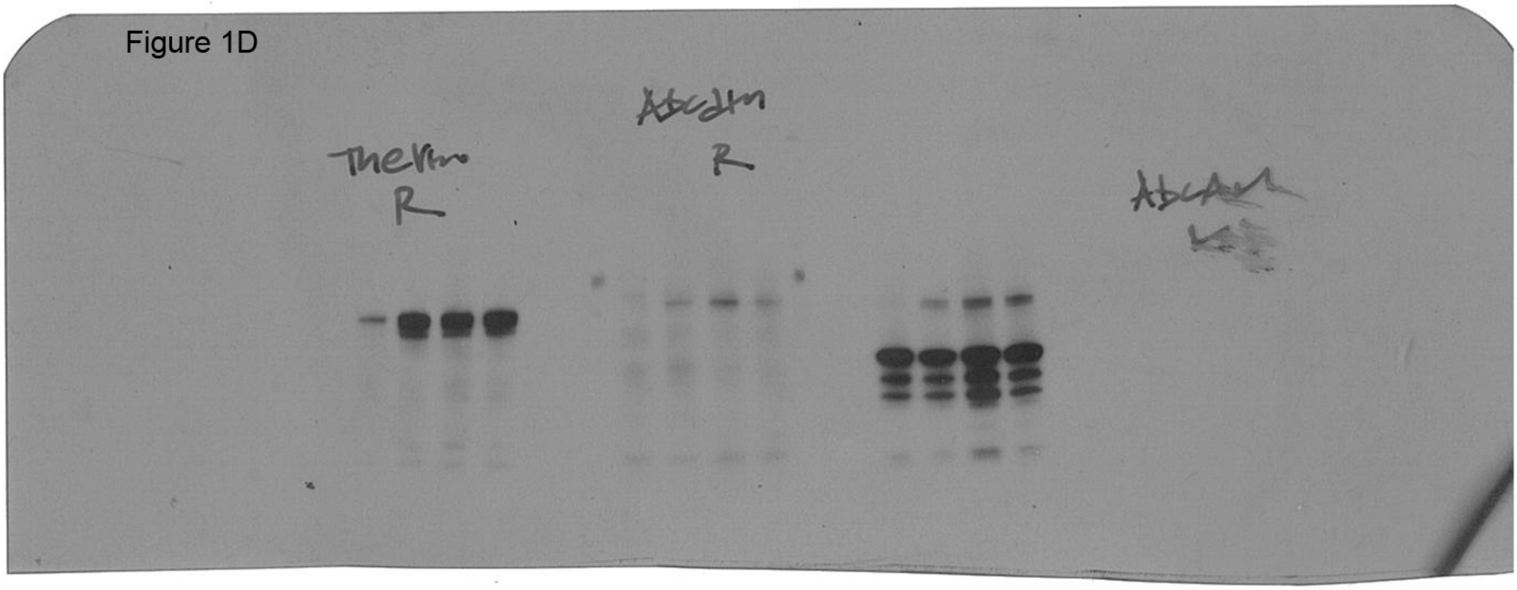

Figure 3I

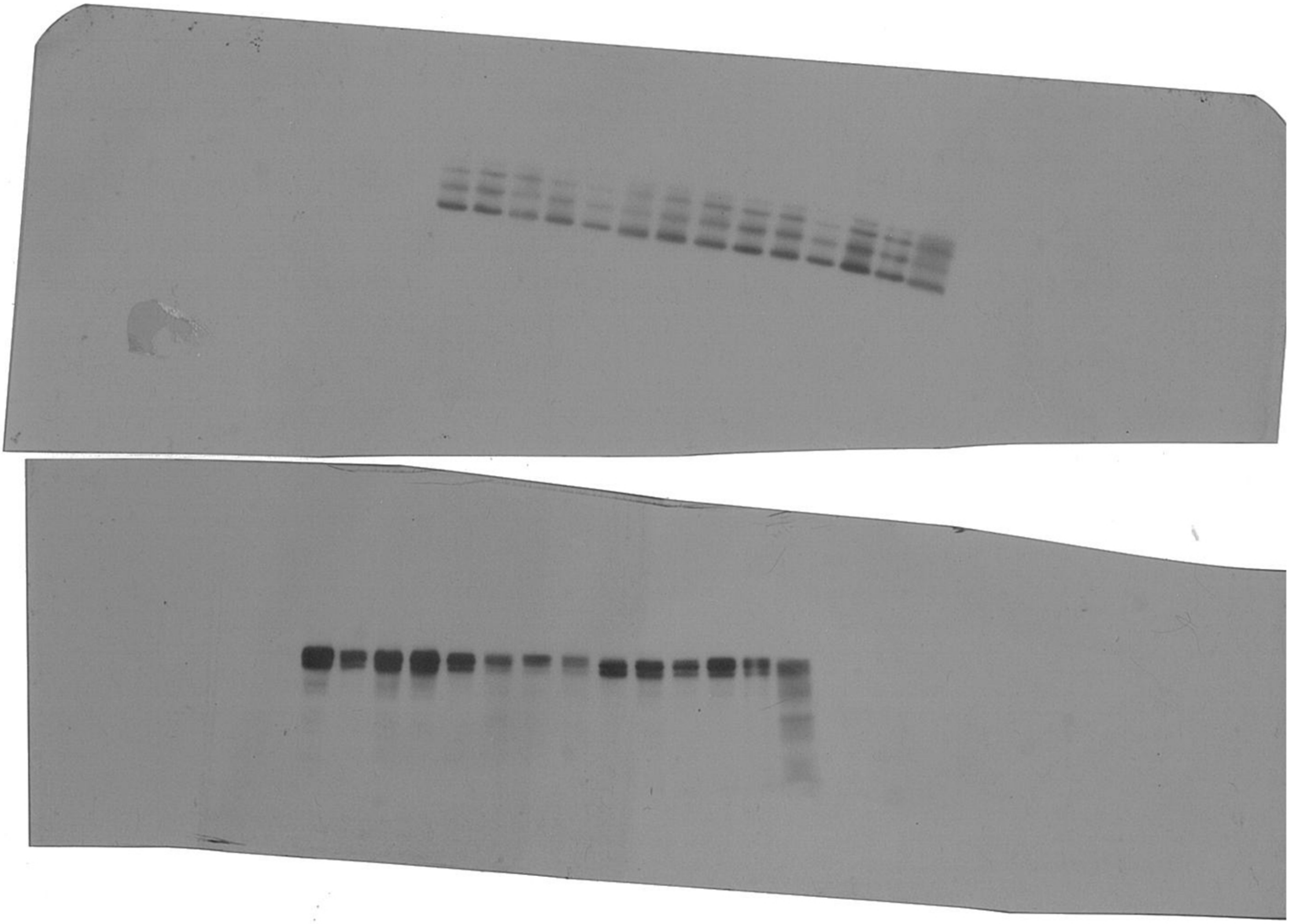

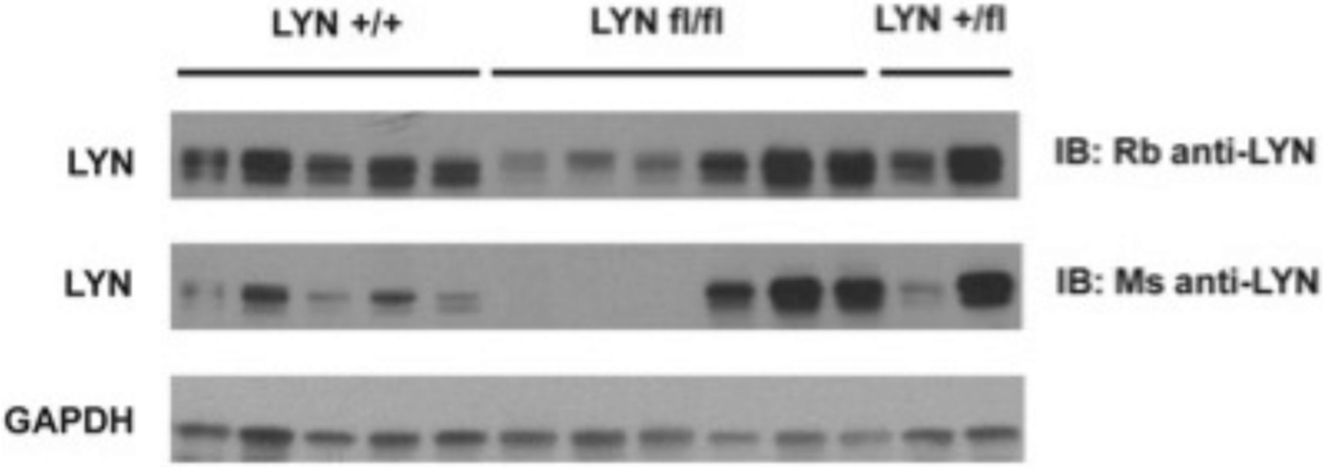

Figure 3I - additional heterozygous tumours

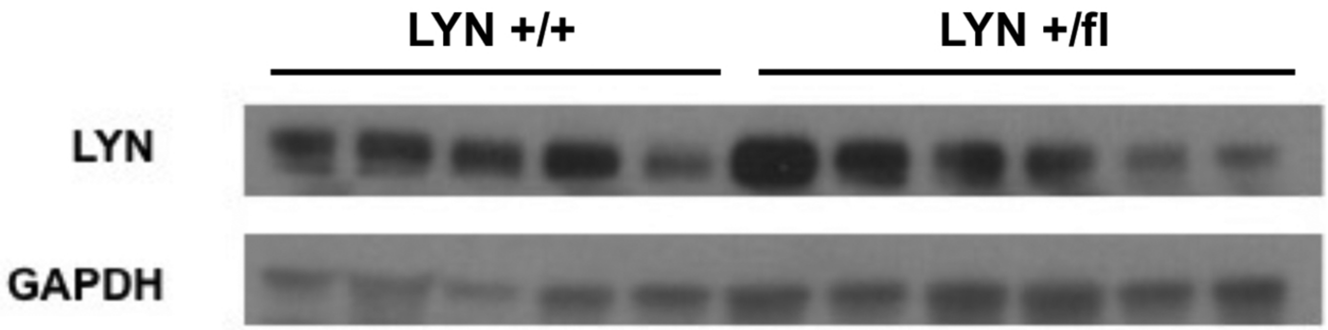


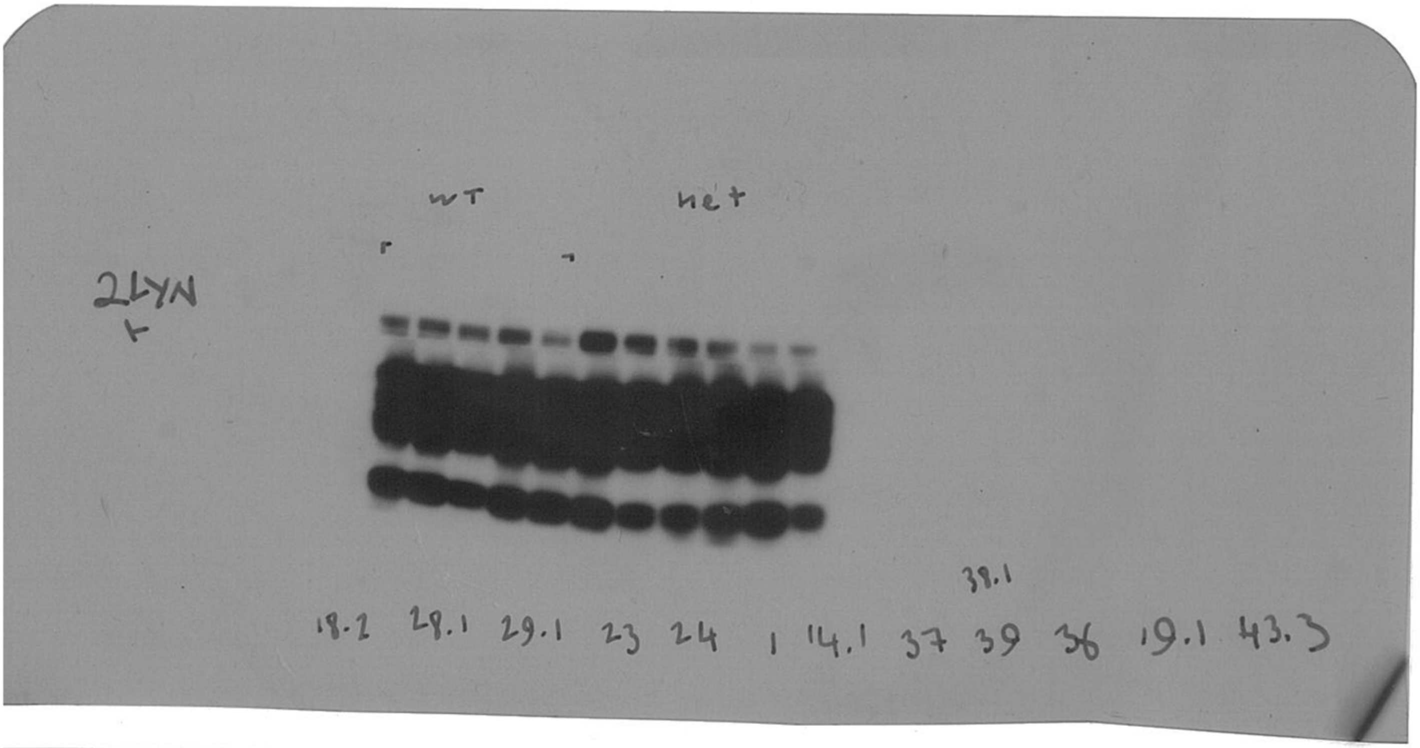


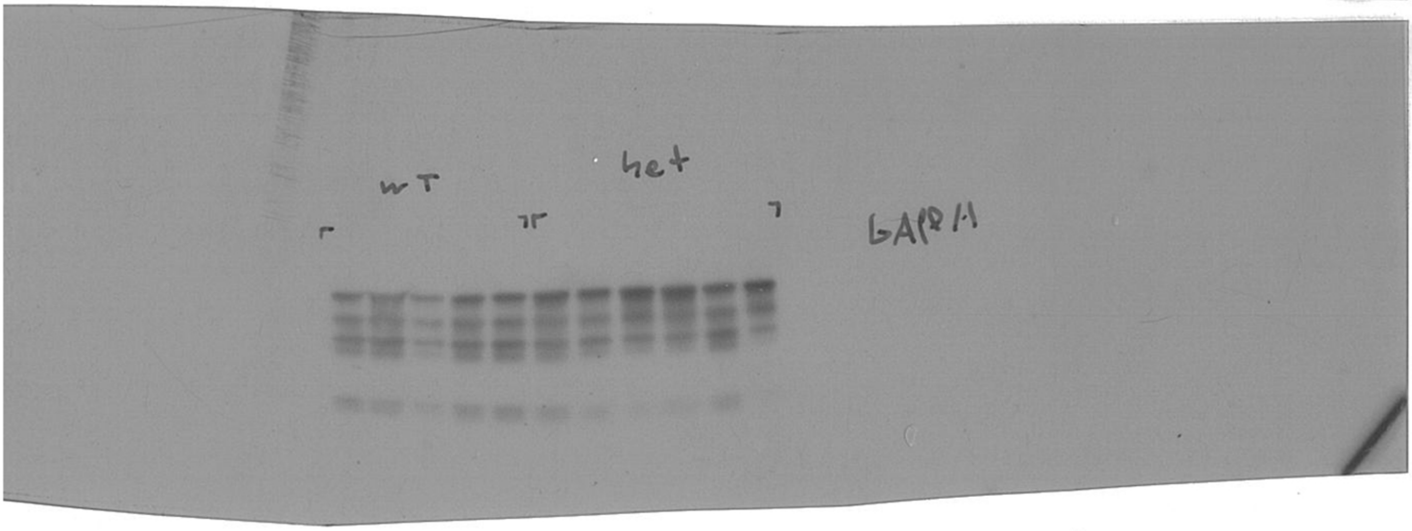
