## Supplementary Legends for "Conditional *in vivo* deletion of LYN kinase has little effect on a *BRCA1* loss-of-function-associated mammary tumour model"

**Supplementary Information**

**Supplementary Figures**

**Each figure is placed on the page above the related legend.**

**Figure S1 (relates to Methods):** Overview of mouse breeding strategy designed to generate desired experimental genotypes (indicated by genotypes in red) while homogenising, as far as possible, any background strain influence across the experimental cohorts.

**Figure S2 (relates to Figure 2)**: H&E stained examples of non-mammary tumours. **A, B)** Lung metastases. Note squamous metaplasia in (B); this sample was from an animal with a squamous mammary tumour. **C, D)** High and low power views of haemangiosarcoma of spleen. **E, F)** High and low power views of osteosarcoma of rib. **G, H)** High and low power views of osteosarcoma from base of tail. Bars A, B, D, F, H = 200μm. Bars C, E, G = 2mm. Inset in (A) magnified 4x. Inset in (B) magnified 2x.

**Figure S3 (relates to Figure 2):** LYN staining in white pulp of spleen. Representative white pulp areas with germinal centres from spleens of *Lyn^wt/wt^* **(A – C)** and *Lyn^fl/fl^* **(D – F)** mice. Antigens detected by immunohistochemistry are indicated on each panel. LYN expression is downregulated in proliferative germinal centres in both genotypes. Bars = 200 μm.

**Figure S4 (relates to Figure 2):** Key distinguishing features enabling histotyping of mouse mammary epithelial tumours. Left hand column, H&E staining. Right hand column, p63 staining. **A, B)** Adenocarcinoma of no special type (AC) showing no metaplastic features (A) and little p63 staining (B). **C, D)** Adenomyoepithelioma (AME) showing characteristic abundant p63 staining with a pseudo-basal pattern and ‘ectopic’ pseudo-luminal expression. **E, F)** Metaplastic adenosquamous carcinoma (ASQC) showing squamous metaplasia, keratin pearls (asterisks) and abundant p63 staining. **G, H)** Metaplastic spindle cell carcinoma (MSCC) showing spindle cell metaplasia and characteristic background stain at the periphery of each spindle cell. This is seen with all antibodies, not just the p63 stain. Bars = 200 μm. Insets are magnified 2.5 times.

**Figure S5 (relates to Figures 2, 3 and 4):** Details of tumour features. **A)** Mammary epithelial tumour histological subtypes by *Lyn* cohort. AC, adenocarcinoma of no special type; AME, adenomyoepithelioma; ASQC, metaplastic adenosquamous carcinoma; MSCC, metaplastic spindle cell carcinoma. No significant differences by Kruskal-Wallis or individual Mann-Whitney tests (n= 21 tumours from *Lyn^wt/wt^* cohort, 47 tumours from *Lyn^fl/wt^* cohort, 37 tumours from *Lyn^fl/f^*^l^ cohort). **B)** Mean number of mammary tumours per animal (±s.d.) by *Lyn* cohort. No significant differences by ANOVA or individual t-tests. The numbers of mice analysed in each group is indicated above each bar. **C)** Example of LYN staining of a tumour assessed at 0.2x, 2x and 20x magnification using OLYVIA software. Scale bars as indicated. This tumour was scored as a ‘3’ for intensity of staining as the staining was apparent even at the lowest magnification. **D)** Normalised RNAseq *Lyn* counts in tumours from PCA groups 1 (n=15), 2 (n=14), 3 (n=6) and 4 (n=4) presented separately. Mean±s.d. indicated. *P<0.05 (ANOVA). **E)** Histoscore of tumours from PCA groups 1 (n=14), 2 (n=11), 3 (n=4) and 4 (n=4) presented separately. Mean±s.d. indicated. *P<0.01 (Kruskal-Wallis). **F)** *In vivo* doubling time (days) of tumours from PCA groups 1 (n=14), 2 (n=13), 3 (n=5) and 4 (n=4) presented separately. Mean±s.d. indicated. No significant differences (Kruskal-Wallis).

**Figure S6 (relates to Figure 7):** Quantitative real-time rtPCR (qrtPCR) expression (raw Ct values) of *Gapdh* and *B-actin* housekeeping genes in whole tumour samples (as used for RNAseq experiments; n= 11 PCA 1/2 and 9 PCA 3/4 tumours, 8 *Lyn* low and 6 *Lyn* high tumours) and tumour cells (n=6 of each genotype) in primary culture. There are no significant differences between any group (Mann-Whitney), however, there does appear to be a batch effect in the cultured cells, with higher *B-actin* expression in *Lyn^wt/wt^* samples. Therefore only *Gapdh* was used for normalisation in these samples.

**Figure S7 (relates to Figure 8):** CIBERSORTx estimates of immune cell abundance in the *Lyn* low (n=13) vs *Lyn* high (n=13) **(A)** and PCA group 1/2 (n=29) vs PCA group 3/4 (n=10) tumours **(B)** based on normalised RNAseq expression data. Estimates are provided using the absolute abundance (arbitrary units) approach (Newman et al., 2019) so that the proportions of each different immune cell type are comparable. Mean±s.d.. N.S., no significant difference, multiple two-tailed t-tests with Holm-Sidak correction. If no significance indication is given, there were too few samples for statistical comparison (or none). ‘Zero’ data points are not plotted. Raw data in **Supplementary Table 13.**

**Supplementary Tables**

**Table S1:** Antibodies and primer sequences used in this study.

**Table S2:** Full details of tumour cohort animals. For mammary tumour phenotypes: AC, adenocarcinoma, no special type; AME, adenomyoepithelioma; ASQC, adenosquamous carcinoma; MSCC, metaplastic spindle cell carcinoma. For mammary tumour locations, the coding indicates the left or right side of the body, and the mammary gland number, 1 – 5. Where more than one tumour has arisen in the same location, they are indicated as A/B/C.

**Table S3:** Full details of analysis of samples taken at necropsy, including animal and sample number, sample type, gross observations and histological observations, presence of metaplastic elements and ASQC or AME-like p63 staining, diagnosis, and, for the mammary tumours, *in vivo* doubling time and RNAseq sample identification. *’Yes’ or ‘No’ indicates whether or not p63 staining pattern was consistent with a diagnosis of ASQC or AME. **Diagnoses of reactive hyperplasia of the spleen versus lymphoma are proposed by MJS and SB on the basis of the clinical behaviour of the animal and histology of the viscera. Extensive phenotyping of lymphocytic populations and analysis of clonality of expanding T-/B-cell populations was not carried out as this was not the primary purpose of the study.

**Table S4:** Ki67 and LYN staining quantitation. Ki67 staining was determined using the Cognition Master Professional Ki67 Quantifier automated Ki67 counting program on five (in one case six) fields of view from each tumour. LYN staining was indicated qualitatively (Y/N) for presence of stained epithelial-like areas, stained individual cells, stained tumour vasculature and stained non-tumour cells in the connective tissue. A histoscore approach (staining intensity 0 – 3 multiplied by area of positive stained cells 0 – 3) was then used to quantify staining in the epithelial-like tumour cells. See Methods for details.

**Table S5:** RNAseq sample identifiers and sample features, including histological observations, diagnosis, *in vivo* doubling time, LYN and Ki67 staining. Also included are the three categories used to identify sets of differentially expressed genes, namely (1) genotype of the animal from which the tumour was derived; (2) the normalised *Lyn* RNAseq expression score and the ranking of each tumour on the basis of that score; (3) the identifiers of each tumour used in the Principal Component Analysis (PCA ID) and its PCA group.

**Table S6:** Raw and normalised RNAseq counts across all 39 samples and analysis of differential expression comparing tumours from the different cohort genotypes.

**Table S7:** Raw and normalised RNAseq counts from the 13 tumours with the highest *Lyn* RNAseq expression counts (*Lyn* high tumours) and the 13 tumours with the lowest *Lyn* RNAseq expression counts (*Lyn* low tumours) and analysis of differential expression of genes across *Lyn* high and *Lyn* low groups.

**Table S8:** Raw and normalised RNAseq counts and differentially expressed genes comparing tumours from PCA groups 1 and 2 with tumours from PCA groups 3 and 4. Note positive fold changes are elevated in groups 3 and 4; negative fold changes are lowered in groups 3 and 4 / elevated in 1 and 2.

**Table S9:** Significantly differentially expressed genes (DEGs) (adjusted p value <0.05; log2 fold change ≤0.5 or ≥2.0) for the WT vs HOM, *Lyn* high vs *Lyn* low and PCA 1 and 2 vs PCA 3 and 4 comparisons. For each comparison, whether or not the DEG is also found to be differentially expressed in the other comparisons is indicated.

**Table S10:** g:Profiler gene set enrichment analysis (GSEA) of DEGs from the *Lyn* expression and PCA comparisons.

**Table S11:** Summary of GO Bioprocess and KEGG pathway interactions. The GO BP terms have been collected together in functional groupings (see first tab) for each tumour set for ease of interpretation. Each tumour set is listed on a different tab. The KEGG pathways enriched within the tumour sets are summarised together on a single tab. Overlaps between GO BP and KEGG Pathway terms between the tumour groups as determined by VENNY are also indicated.

**Table S12:** Raw and normalised quantitative real time rtPCR data from validation of differential gene expression (**Figure 7** and **Figure S6**). The first tab has raw Ct values, the second tab has values for whole tumour samples (as used for RNAseq) normalised by geometric mean of *Gapdh* and *B-actin*, the third tab has values for cultured cells normalised by *Gapdh* only. Normalised values indicated in grey, failed wells in red, samples used as comparator populations in yellow.

**Table S13:** Results of CIBERsortx analysis on *Lyn* expression and PCA tumour groups.

**Supplementary Data**

**Supplementary Data File:** Raw scanned western blots from Figure 1C, D and Figure 3.
